# Perceptual glimpses are locally accumulated and globally maintained at distinct processing levels

**DOI:** 10.1101/2025.04.30.651428

**Authors:** Elisabeth Parés-Pujolràs, Anna C. Geuzebroek, Redmond G. O’Connell, Simon P. Kelly

**Affiliations:** School of Electrical and Electronic Engineering and UCD Centre for Biomedical Engineering, University College Dublin, Dublin, Ireland; Trinity College Institute of Neuroscience and School of Psychology, Trinity College Dublin, Dublin, Ireland

## Abstract

Making decisions often requires the integration of multiple pieces of information. An extensive body of research has investigated the neural architecture underpinning evidence accumulation in perceptual tasks where information is continuously present, but less is known about how this neural architecture operates in situations affording only intermittent glimpses of an evidence source. In two electroencephalography (EEG) experiments, participants judged the direction of up to two pulses of motion evidence separated by gaps of varying duration. Our behavioural analysis found that participants used both pulses but underutilised the second, and showed no systematic decrease in accuracy as a function of gap duration. At the neural level, motor beta lateralisation tracked cumulative evidence across pulses, maintaining a sustained representation of the decision variable through the gap and until response. In contrast, the centroparietal positivity (CPP), a previously-characterised signature of evidence accumulation, built up transiently to a peak that scaled with each pulse’s contribution to the decision variable (i.e. the absolute belief update it produced), falling back to baseline in between pulses. These patterns were recapitulated in a model where pulse-information transiently integrated at the CPP level is fed to and maintained at a bounded motor level. In this model, the evidence in the second pulse is only integrated to the extent that the evidence in the first pulse falls short of the bound, or not integrated at all if a bound has already been hit.

## 1. Introduction

Perceptual decision-making is widely thought to involve a process of evidence integration over time (Hyafil et al., 2023). An extensive body of research has successfully linked these integration computations to multiple neural decision signals in various species (Gold & Shadlen, 2007; Hanks & Summerfield, 2017; Heekeren et al., 2008; O’Connell & Kelly, 2021; Shadlen & Kiani, 2013) revealing a neural decision architecture that includes both effector-selective motor preparation signals, and motor-independent signals that appear to reflect integration operations at an intermediate processing level (Gherman et al., 2024; O’Connell & Kelly, 2021; Okazawa & Kiani, 2023; Stockart et al., 2025). Such neural decision processes have mainly been studied in the context of tasks requiring immediate judgements about a single continuously-presented stimulus. However, in many real-life situations a decision-maker is afforded only a short glimpse of a relevant evidence source and it is uncertain whether and when they will receive an additional glimpse, as, for example, when evaluating an object among moving traffic that may be intermittently occluded. This scenario has begun to be studied using an intermittent-evidence task (Kiani et al., 2013), which we refer to as “gaps task”, in which up to two short pulses of noisy evidence for a certain motion direction are presented separated by a variable temporal gap, with a behavioural response required after a delayed cue. Previous studies using this task or variants of it have found that both evidence pulses are utilised, but not fully or equally (Azizi & Ebrahimpour, 2023; Golmohamadian et al., 2025; Kiani et al., 2013; Tohidi-Moghaddam et al., 2019), and although a general information leakage account could be ruled out based on the absence of any decline in accuracy with gap duration, the neural underpinnings of the task, including the functional roles of effector-selective and motor-independent processes in handling and maintaining the intermittent evidence, and how such information underutilisation can arise, have remained unclear.

In humans, two neurophysiological signatures of decision formation can be utilised alongside behavioural data to gain insight into mechanisms underlying perceptual tasks (O’Connell & Kelly, 2021). First, spectral activity in the beta band (13-30 Hz), long observed to decrease over motor cortex with preparation of a contralateral limb (Pfurtscheller & Lopes da Silva, 1999; Stancák & Pfurtscheller, 1995), has been found in decision tasks to build at an evidence-dependent rate, reaching a threshold level contralateral to, and coincident with, the executed action (Donner et al., 2009; Feuerriegel et al., 2021; Geuzebroek et al., 2023; Kelly et al., 2021; O’Connell et al., 2012; Steinemann et al., 2018). Motor beta dynamics have also been shown to reflect decisions and choice biases in humans (de Lange et al., 2013; Gould et al., 2012; Herding et al., 2016; Pape & Siegel, 2016; Urai & Donner, 2022; Wyart et al., 2015) and non-human primates (Haegens et al., 2011, 2017; Rassi et al., 2023), highlighting its role as a key marker of decision-related activity. Second, a centroparietal positivity (CPP) in the human electroencephalogram (EEG) has been found to exhibit similar evidence-dependent buildup dynamics in conventional perceptual tasks with continuous stimuli such as random dot motion, but with an earlier latency; it begins building at a stimulus-aligned onset time, rises at a rate that scales with evidence strength, and peaks at the time of action choice when immediate execution is allowed (Kelly & O’Connell, 2013; O’Connell et al., 2012; Steinemann et al., 2018). Signals with similar dynamics have been observed in multiple brain regions including the parietal cortex intracranial recordings (Gherman et al., 2024; Pereira et al., 2021; Stockart et al., 2025), bearing some overlap with regions identified in functional imaging studies (Liu & Pleskac, 2011; Ploran et al., 2007; Tremel & Wheeler, 2015). In contrast with motor preparation signals, the CPP exhibits evidence-integration dynamics even when no motor output is required (O’Connell et al., 2012). In delayed-response tasks where actions are deferred until some time after the perceptual choice is made, the CPP peaks at decision commitment and falls back to its baseline level (Azizi & Ebrahimpour, 2023; McCone et al., 2026; Twomey et al., 2016), whereas relative motor preparation reflected in motor beta lateralisation (MBL) sustains until action execution (Donner et al., 2009; McCone et al., 2026; Murphy et al., 2021; Parés-Pujolràs et al., 2025; Twomey et al., 2016). These findings are consistent with the CPP tracking an intermediary, motor-independent evidence integration process that is continuously fed to downstream motor preparation, where a sustained representation of the DV is maintained after commitment is reached, until a response is made.

More recently, a few studies have highlighted that the CPP can behave differently in expanded judgement tasks, where evidence is presented as a series of discrete tokens that need to be integrated rather than as a continuous, noisy stream. In these tasks, centroparietal signals exhibit transient token-evoked positivities that scale with the immediate evidence, rather than tracking the evolution of an ongoing decision variable (DV) across tokens (Parés-Pujolràs et al., 2025; Wyart et al., 2012, 2015). Motor Beta Lateralisation (MBL), meanwhile, tracks the evolving decision variable across discrete tokens over multiple seconds, by stepping up or down according to the evidence in each token and sustaining in between them (Murphy et al., 2021; Parés-Pujolràs et al., 2025; Wilming et al., 2020). These findings again point to the operation of an intermediary decision process reflected in the CPP - one that can be flexibly configured to compute different quantities depending on the nature of the task.

The Gaps task, like expanded judgment tasks, involves temporally disparate pieces of information, but it holds key distinctions that demand dedicated investigation. In expanded judgement tasks, tokens typically bear little or no perceptual noise, but the overlap of the underlying generative distributions means they often provide conflicting information (e.g. Balsdon et al., 2020, 2021; Glickman et al., 2022; Rouault et al., 2022; Waskom & Kiani, 2018; Wyart et al., 2015). The nature of the stimuli thus encourages appraisal of each token as a distinct object, potentially explaining the transient pattern of CPP activity. In contrast, the Gaps task involves pulses of evidence that are perceptually noisy but always indicate a consistent motion direction (Kiani et al., 2013), thus appearing as glimpses of the same object rather than a series of individual objects. Only one study has examined neural signatures of decision formation in this intermittent evidence context: using EEG, Azizi & Ebrahimpour (2023) showed that the CPP exhibits an evoked positivity after each pulse that appears to decay during the gap, suggesting a local processing role. However, how these pulse-evoked activities relate to the computations underpinning an ongoing decision, interact with motor preparation to determine choice, and relate to any information underutilisation, has not been examined.

In this study, we recorded EEG during the Gaps task (Kiani et al., 2013) in two experiments with slightly different settings for the variable temporal gaps to ensure generality of findings. Our primary goal was to characterise the dynamics of the CPP and MBL in the context of this intermittent evidence task, and functionally relate them to behavioural indices of integration across tokens. We found that participants based their choices on both pulses but with a weaker influence of pulse 2, and coherence-dependent weightings on choice were mirrored in the magnitude of transient pulse-evoked CPPs. These patterns could be explained by a decision architecture where the CPP conducts two separate rounds of evidence accumulation, continuously feeding the accumulating evidence within each pulse to a downstream motor process indexed by MBL. MBL in turn tracks a continuous representation of the state of the overall DV, sustaining without loss of information throughout the gap and between the final evidence and the response. Our behavioural and modelling results indicated that participants engaged in a bounded accumulation process, so that evidence samples within each pulse were only integrated while the DV encoded in MBL had not yet reached the bound. Additionally, given the dynamic structure of the task involving evidence onsets and offsets, we explored the dynamics of attention, as indexed by occipital alpha signals, and these were suggestive of a key role in filtering-out of irrelevant dot-motion, and in modulating the weight of pulses on choice.

## 2. Results

### 2.1. Behavioural results

In this study, participants engaged in a motion direction discrimination task where, on most trials, two pulses of partially-coherent dot motion (indicated by yellow dot colour) were separated by variable temporal ‘gaps’ in which motion was random, and thus uninformative (indicated by blue dot colour; *Fig. 1A*). Though coherence could differ, the direction was always consistent between the two pulses, so that they would be construed as two glimpses of the same evidence source. Our two experiments (Exp. 1: N = 22; Exp. 2: N = 21) differed only in the delays separating the two pulses. In both experiments, coherence levels were titrated for each participant using a staircase procedure aiming for an average 70% accuracy following a single evidence pulse. High (Exp. 1: M = 44.73%, SD = 12.64%; Exp 2: M = 33.04%, SD = 13.41%) and low (Exp. 1: M = 26.75%, SD = 7.74%; Exp 2: M =19.82%, SD = 8.04%) coherences were then set to be, respectively, 25% higher and 25% lower than the staircase-estimated coherence (see *Methods*), and small adjustments were made between blocks if performance substantially changed (*Figure S1*).

**Figure 1.**
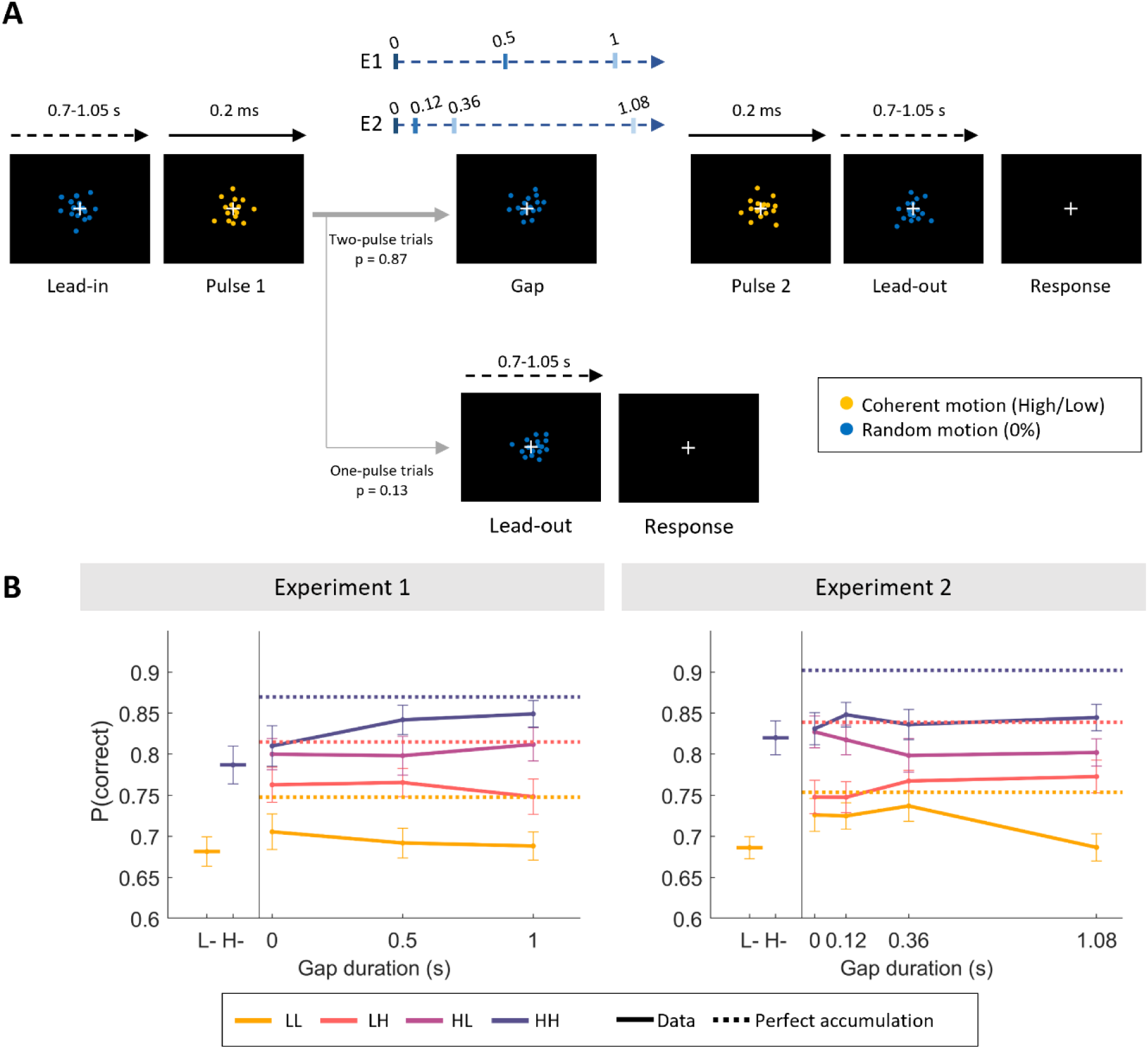
Task and behavioural results. ***A***. Experimental structure. Participants viewed a random dot stimulus that coherently moved either to the left or right during yellow evidence pulses, of which there was sometimes one, but usually two, separated by a variable gap in evidence indicated by blue dot colour. **B.** Grand-averaged (mean ± SEM) accuracy in two-pulse trials, sorted by pulse coherence (low, L/ high, H), order (LL, LH, HL, HH) and gap duration. Data points in the left-hand section indicate single-pulse accuracies. Overlaid horizontal dashed lines indicate the expected accuracy of a perfect accumulator, based on the grand-average performance on one-pulse trials. Note that a perfect accumulator predicts identical accuracies for the HL and LH conditions, and therefore the two lines overlap.

As expected, participants’ accuracy scaled with motion coherence and number of pulses. In one-pulse trials, participants were more accurate in the high coherence compared to the low coherence trials (*Eq. 1*, Exp 1: *β_1_* = 0.33 ± 0.05, *p <* 0.001; Exp 2: *β_1_* = 0.43 ± 0.06, *p <* 0.001, *Figure 1A*). There was a further benefit in accuracy from the second pulse when present, indicated by sensitivity to coherence in both the first (*Eq.2,* Exp 1: *β_1_* = 0.26 ± 0.06, *p <* 0.001; Exp 2: *β_1_* = 0.31 ± 0.05, *p <* 0.001) and second (*Eq.2,* Exp 1: *β_2_* = 0.16 ± 0.06, *p =* 0.001; Exp 2: *β_2_* = 0.06 ± 0.04, *p =* 0.015) pulses. When both pulses in a two-pulse trial were the same coherence, participants performed better compared to one-pulse trials, exhibited both in high and low coherence trials in Experiment 2 (*Eq. 3*, Exp. 2: *β_1_* = 0.16 ± 0.08, *p =* 0.047*; β_3_* = 0.02 ± 0.08, *p =* 0.784), but only significantly so in high coherence trials in Experiment 1 (*Eq. 3*, Exp. 1: *β_1_* = 0.21 ± 0.07, *p =* 0.013*; β_3_* = 0.16 ± 0.07, *p =* 0.019). However, they did not perform as well as expected from a perfect accumulator (*Figure 1B*), estimated using performance in one-pulse trials to predict expected accuracy in two-pulse ones (see *Eq*. 5-7; see *Methods*).

The fact that the regression coefficients for the second pulse were smaller than those estimated for the first one indicates that participants’ choices were guided more strongly by the first piece of evidence they received. On average, accuracies were lowest for the most difficult trials (both pulses had low motion coherence, LL), intermediate for trials with mixed coherences (LH, HL) and highest for the easiest trials (both pulses had high motion coherence, HH). However, we also observed a significant pulse-order effect: when the two presented pulses had different coherences, participants were more accurate when the first pulse was the high-coherence one (HL) compared to trials where the first pulse was low-coherence (LH) in both experiments (*Eq. 4*, Exp 1: *β_2_* = -0.11 ± 0.03, *p* <0.001; Exp 2: *β_2_* = -0.12 ± 0.02, *p* <0.001; *Figure 1B)*. This is in line with participants’ choices being more strongly influenced by the first evidence pulse.

Importantly, there was no main effect of gap duration on accuracies (*Eq. 2*, Exp 1: *β_3_* = 0.06 ± 0.08, *p =*0.525; Exp 2: *β_3_* = -0.003 ± 0.05, p =0.579; Figure 1B), suggesting an absence of leakage of information from the first pulse, in line with previous work (Azizi & Ebrahimpour, 2023; Kiani et al., 2013). However, in Experiment 2, gap duration interacted with the coherence of the second pulse (*Eq 2*, Exp 2: *β_5_* (p2 x gap interaction) = 0.16 ± 0.07, *p =*0.031). Since this effect was inconsistent across experiments and we did not have prior hypotheses about it, we leave it to one side and focus on the main effects of coherence. In what follows, we combine EEG and computational modelling to test the possibility that the greater sensitivity to the first pulse arises from the participants engaging in a bounded accumulation process where decisions are sometimes terminated early, curtailing the extent to which the second pulse benefits accuracy and driving the coherence order effects.

### 2.2. EEG results

#### 2.2.1. Activity during the gap

We then turned to neural correlates of evidence accumulation during the task, specifically, MBL and the CPP. Across the two experiments and in all gap conditions, MBL showed evidence-dependent buildup continuously through the sequence of pulses with a temporal lag (*Fig. 2A*), sustaining at a coherence-dependent level (see *Fig. S2; S3*) throughout the gap between the two pulses and until response, showing stronger lateralisation towards the correct response following high-coherence pulses. In contrast, the CPP exhibited a transient build-up following each evidence pulse, falling back to baseline levels prior to the second pulse (*Fig. 2B)*.

**Figure 2.**
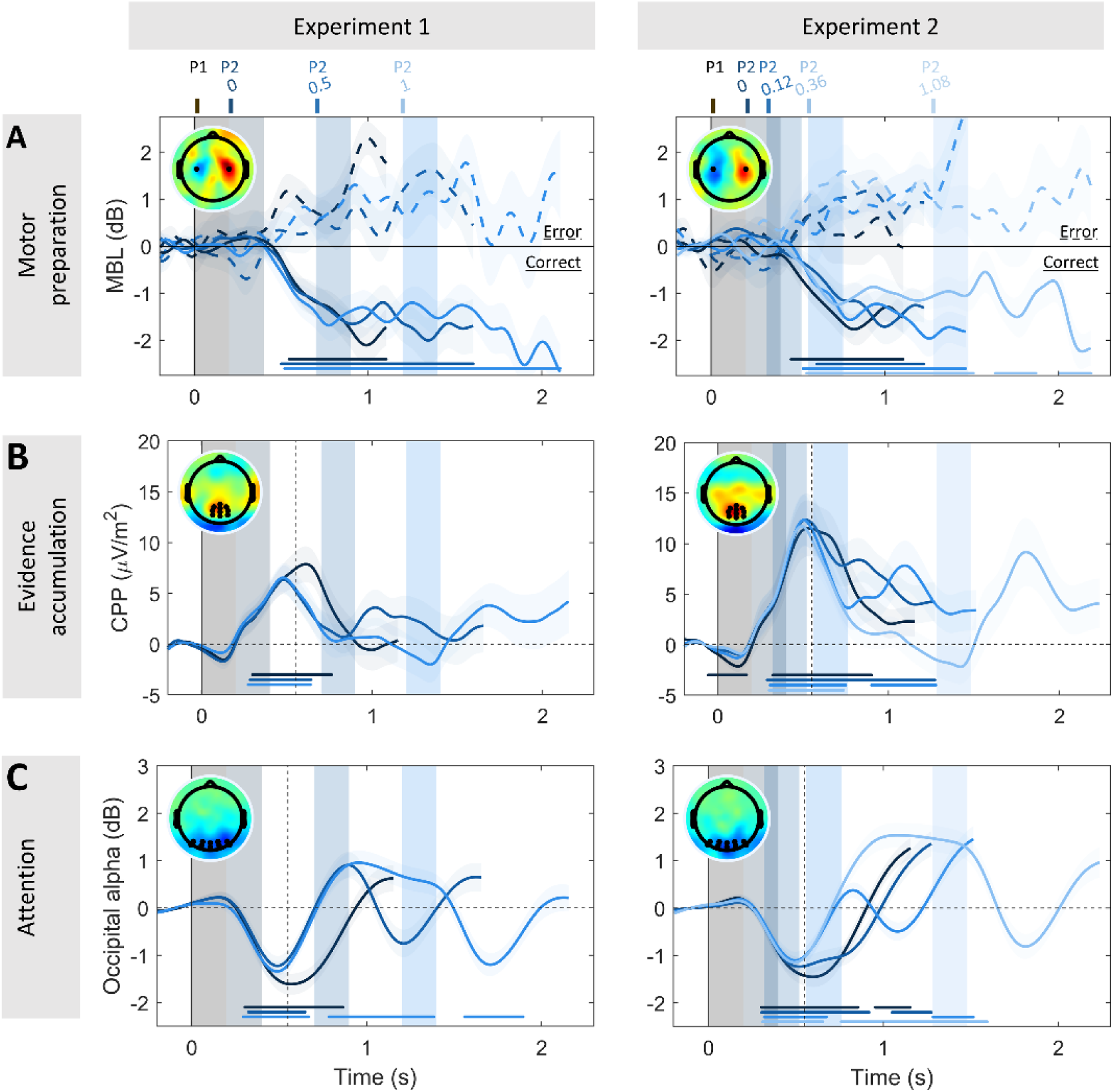
Neural correlates of evidence accumulation and motor preparation across temporal gaps. **A.** Grand-averaged (±SEM) motor beta lateralisation (MBL) [Contra-Ipsilateral hemispheres with respect to correct response hand], sorted by accuracy (*solid line* correct, *dashed line* error). Participants started preparing a motor response in response to the first pulse, and MBL was sustained throughout the gap and until the time of behavioural response across all gap conditions in both experiments. **B.** Grand-averaged (±SEM) centroparietal positivity (CPP) across all temporal gap conditions. The CPP showed transient evoked responses following each pulse. Note that the evoked amplitudes by P1 (CPP-P1) were systematically higher than those evoked by P2 (CPP-P2) (Exp. 1: CPP-P2 = 52% of CPP-P1 peak amplitude; Exp. 2: CPP-P2 = 58% of CPP-P1 peak amplitude; see also *Fig. S8*). **C.** Grand-averaged (±SEM) occipital alpha power for each of the gap conditions, baseline-corrected before the onset of the first pulse and collapsed across coherence. Occipital alpha power showed a steep decrease following pulse onset (which was clearly indicated by a change in the colour of the moving dots), and steep increase overshooting the baseline level once the dots changed colour again to indicate non-coherent motion. Note that for both motor beta and alpha, greater negative values reflect greater preparation of the correct response and attention, respectively. In all panels, shaded areas indicate the timing of evidence pulses for the various gap durations, which are colour coded (see top axis). Topographies illustrate MBL before response (right minus left responses, ***A***) in correct trials, and CPP/alpha power 550 ms post stimulus onset (*dashed line, **B,C***). Further, markers along the bottom indicate times where the signal significantly differed from zero (two-tailed cluster-based permutation test, p<0.005) in only correct trials for MBL for consistency of polarity (***A***), and for all trials pooled in the case of the CPP (***B***) and alpha (***C***). Note that evidence-dependent buildup dynamics are clearly delayed in MBL relative to the CPP, and this aligns with previous research supporting the contention that the CPP lies upstream from motor preparation (Steinemann et al. 2018); such time delays were not a focus of the present study because in such delayed-response conditions we do not have access to the time of decision commitment.

Given that our task included strong external (colour) cues indicating time windows with and without evidence, we further investigated the dynamics of attentional engagement with the stimulus, as indexed by occipital alpha activity (Foxe & Snyder, 2011; Van Diepen et al., 2019), throughout trial time. In both experiments, we observed strong occipital alpha desynchronisation (i.e. a decrease relative to baseline) during the periods in which the CPP indicated active evaluation of the yellow-dot evidence periods (*Fig. 2C*), and strongly elevated alpha above baseline levels in between those pulse-processing periods. Adding to the above behavioural finding of no main effect of gap duration, this suggests that the participants successfully gated stimulus evaluation according to dot colour, selectively attending to the temporal windows in which decision-relevant information is available for evaluation, and ignoring the decision-irrelevant gap periods.

#### 2.2.2. Centroparietal signals transiently reflect local accumulation of evidence in each pulse

Next, we examined coherence-dependent CPP dynamics during the pulses. The CPP evoked after the first pulse (CPP-P1) scaled with the strength of evidence as expected, building more steeply (Exp. 1: t_(21)_ = 2.18, *p* = 0.04; Exp. 2: t_(20)_= 3.48, *p* = 0.002) and reaching higher amplitudes after high coherence pulses compared to low-coherence ones over a cluster of centroparietal electrodes (Exp. 1: *p* = 0.009, Exp. 2: *p =* 0.031; two-tailed cluster-based permutation test; *Fig. 3A-D*). Furthermore, single-trial variability in CPP-P1 amplitude significantly modulated the first pulse’s weight on choice: in trials where evoked CPP-P1 amplitudes were particularly high, the coherence of pulse 1 had a stronger influence on behaviour (*Fig. S11A*).

**Figure 3.**
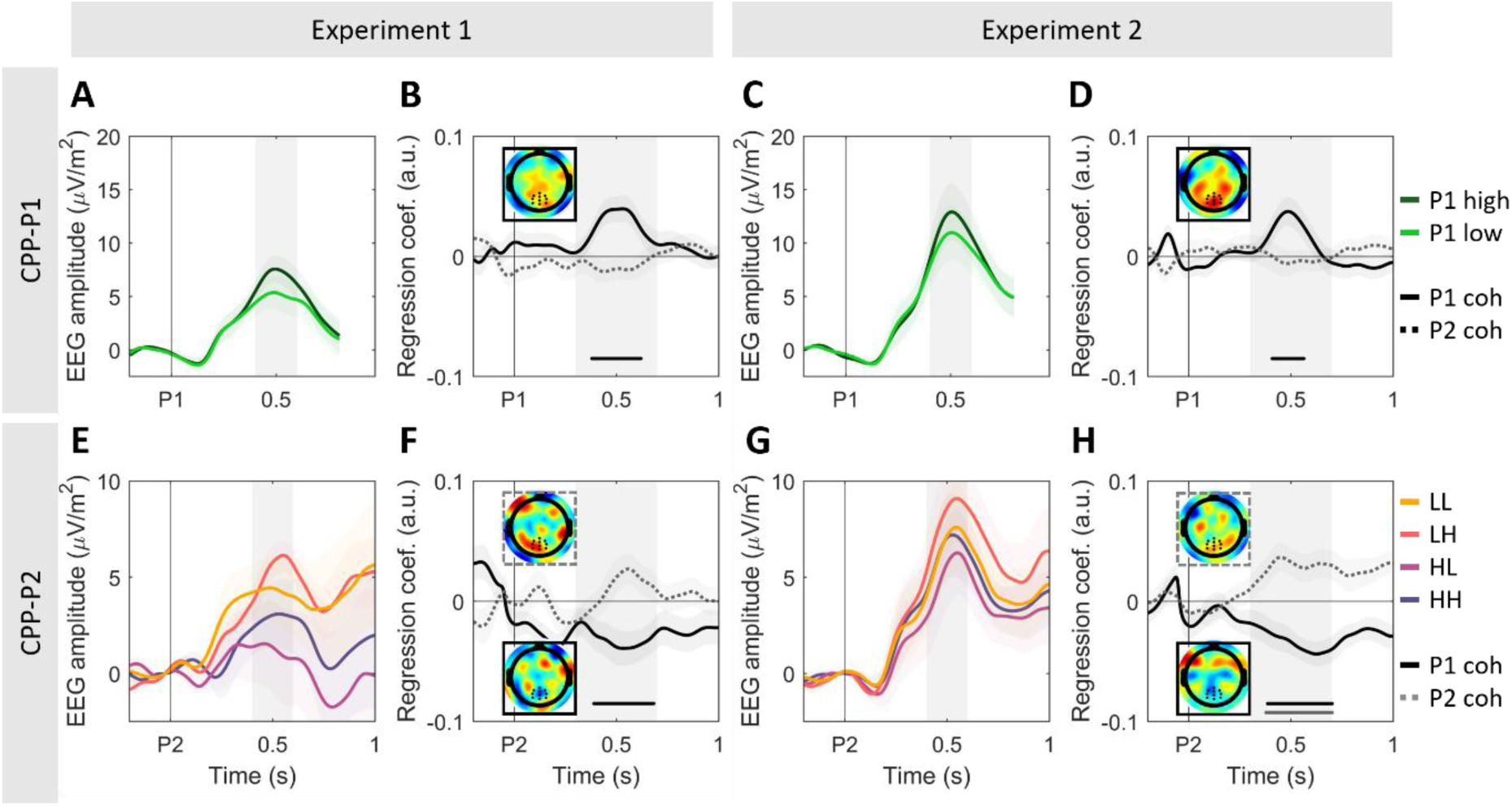
CPP amplitude following each pulse scales with its coherence, and the strength of evidence in pulse 1 additionally inversely modulates amplitudes following pulse 2. **A,C.** Grand-averaged (±SEM) centroparietal activity after pulse 1 (CPP-P1), averaged across all gap conditions. **B,D.** Standardized regression coefficients illustrating the effect of P1 (*solid line*) and P2 (*dashed line*) coherence on evoked CPP-P1 amplitudes. Inset topographies illustrate the coherence effect [P1 high – P1 low] at [0.4-0.6s] after P1. **E,G.** Grand-averaged (±SEM) centroparietal activity after pulse 2 (CPP-P2), averaged over all non-zero gaps in the two experiments. For illustration purposes, we subtracted the average activity of long gap trials with the same P1 coherence from the epoched EEG traces, between [-0.2 to 1.4s] from P1 presentation, and baseline corrected traces [-100 to 0ms] before P2. This isolates the neural activity uniquely associated with the presentation of the second pulse in short gap trials (<1s), where potentials evoked by the first and second pulses overlapped, by removing the activity associated with the first pulse from the P2-evoked traces, including baseline trends (see *Fig. S4*). The qualitative patterns did not vary if this correction was not applied (*see Fig S5*). Note also the difference in the y axis scale compared to panels A,C. **F,H.** Standardized regression coefficients (mean ±SEM) illustrating the effect of P1 (*solid line*) and P2 (*dashed line*) coherence on evoked CPP-P2 amplitudes. Inset topographies illustrate the difference between [P1 high – P1 low], solid outline; [P2 high – P2 low], dashed outline) [0.4-0.6s] after P2. The negative effect of P1 illustrates higher amplitudes for P1-low trials. *Markers along the bottom of panels B,D,F and H indicate significant clusters where coherence effects differed from zero (two-tailed cluster-based permutation test, p<0.05). Shaded areas indicate the tested time window. See *Fig. S3* for full-trial ERP traces, sorted by gap and coherence conditions.

The coherence of the second pulse modulated peak amplitude in the standard way, with larger CPP amplitudes in trials with high pulse 2 coherence, although this effect was only significant in Experiment 2 (Exp. 1: *p =* 0.165, Exp.2: *p =* 0.009; two-tailed cluster-based permutation test; *Fig. 3F,H;)*. CPP-P2 amplitude was also influenced by the coherence of Pulse 1: on trials where the first pulse was low coherence, CPPs evoked by the second pulse reached higher amplitudes (Exp. 1: *p =* 0.018, Exp. 2: *p =* 0.005; two-tailed cluster-based permutation test; see *Fig. 3F,H*). Additionally, CPP-P2 amplitude was generally smaller than CPP-P1 amplitude, which was the case whether we computed CPP-P2 amplitudes after baseline-correction before pulse 2 onset (Exp. 1: t(21) = 2.29, *p* = 0.032; Exp. 2: t(20) = 4.19, p < 0.001) or after correction for CPP-P1 overlap (Exp. 1: t(21) = 1.93, *p* = 0.067 ; Exp. 2: t(20) = 3.25, *p* = 0.003; see *Methods* for details).

#### 2.2.4. A single bounded decision process can account for key behavioural and neural patterns

We then aimed to identify a computational model that could account for the key behavioural accuracy and CPP amplitude effects observed in our data. Similar to previous studies (Azizi & Ebrahimpour, 2023; Kiani et al., 2013; Tohidi-Moghaddam et al., 2019) there was no evidence of leak in our data, given that no general decrease in accuracy was observed for longer gap durations. This lack of a main effect of gap duration on accuracies also indicates that participants did not accumulate the incoherent motion presented during the gaps. Instead, the primacy effect observed in our behavioural results, with higher accuracies in HL than LH trials, is consistent with a bounded process where participants sometimes terminated evidence accumulation early, failing to use all available evidence. Thus, we fitted a single bounded accumulator model with three free parameters (bound, high- and low-coherence drift rates), where evidence accumulation without leak proceeds only during the pulse time windows and is terminated upon bound crossing (see *Methods* for full details). The overall DV was sustained at the value attained at the end of pulse 1 throughout the gap. If the bound was not crossed by the end of the trial, the sign of the DV at the end of the trial determined the model’s predicted choice.

This simple single-DV model could account for the order effects observed in behavioural data, where the first pulse had a greater bearing on choice than the second one, while at the same time capturing the generally greater accuracy of two-pulse trials compared to single-pulse ones (*Fig. 4A*). In this model, this partial benefit of the second pulse arises from the fact that the DV is further influenced by it only on the trials where the DV has not already reached a bound. The second pulse can thus be integrated to varying extents along a continuum: not at all (if a bound is hit during pulse 1), partially (if no bound was hit during pulse 1, and only until a bound is hit during pulse 2), or fully (if no bound is hit at any point during the trial). As expected, a greater proportion of trials terminate early in the P1-high trials compared to P1-low trials (*Fig. 4B*). Note that this simple diffusion model has no way to produce systematic gap-dependent effects, meaning that predicted accuracies are forced to be flat across gap durations; again, since the observed gap-dependent effects were unexpected and differed in our two experiments, we did not pursue further modelling approaches to attempt to capture them in the current study (but see *Fig. S9, S10* and *S11* for exploratory analyses on how temporal expectations might influence integration processes and evidence weighting). We did explore alternative flexible weighting models in which the underutilisation of P2 occurred as a direct effect of dampening the evidence therein as opposed to indirectly through the application of the bound. While these could explain the observed primacy effects, the simple bounded model with no drift rate re-weighting provided the best fit to the current data when accounting for parsimony according to Bayes Information Criterion (see Supplementary Note 1 for an extended discussion).

**Figure 4.**
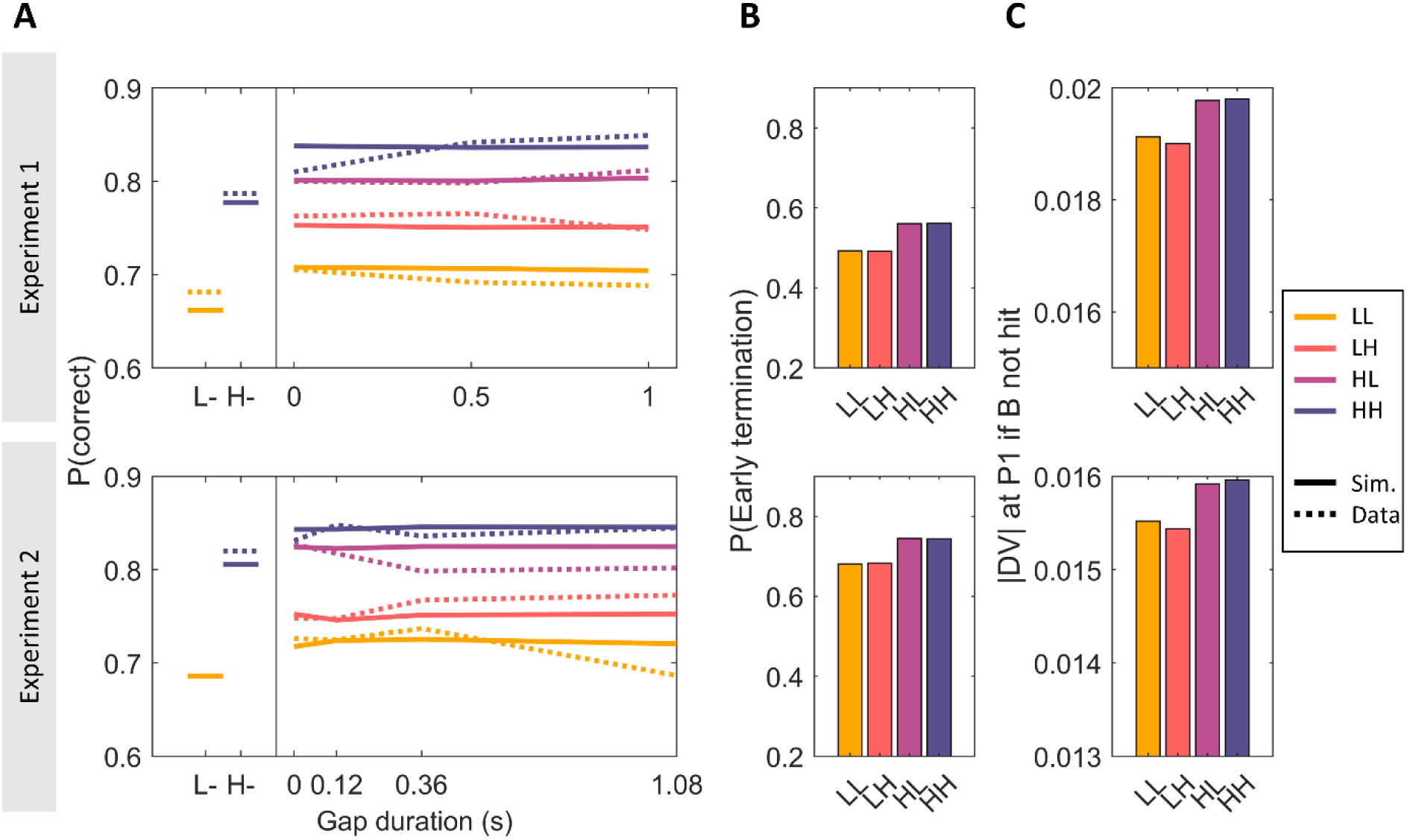
Simulated behavioural data. (N = 50000) of a simple bounded accumulator model with three parameters (B (bound), d1 (low drift rate), d2 (high drift rate)), fit to all grand-averaged data points for each experiment separately (see Fig. S13 & Fig. S14 for results based on single-participant fits). **A.** Grand-averaged simulated (*solid lines*) accuracy in single-(*left*) and two-pulse (*right*) trials, sorted by pulse coherence and gap duration in experiments 1 (*top*) and 2 (*bottom*). Observed data are overlaid (*dashed*) for comparison. Simulated data recapitulated the observed order effects. For simplicity, the model was not endowed with any means to account for effects of gap duration. **B.** Proportion of simulated two-pulse trials that hit a bound during the first pulse, and therefore did not use the second pulse evidence to make a choice. **C.** Absolute value of the decision variable (DV) at the end of pulse 1 in the subset of trials where the bound was not hit during that pulse, sorted by condition.

Then, we simulated, from this diffusion model fit to accuracy data, the neural activity corresponding to MBL and the CPP (*Fig. 5*). Our empirical results suggested that motor preparation as indexed by MBL tracks the state of the bounded decision variable in a sustained manner, emerging shortly after the first pulse and remaining lateralised until response. Thus, we simulated MBL by taking the signed values of the model-derived DV throughout the trial and averaging across trials. These exhibited a similar pattern to the empirical data, with the signed DV showing opposite-sign patterns for error vs. correct trials (*Fig. 5A,D*, *top; cf. Fig. 2A*). If a bound was hit, the DV was set to stay stable in our model, and therefore recapitulated the way in which motor preparation responses here (*Fig 2A*) and in previous studies (e.g. (Donner et al., 2009; Murphy et al., 2021; Parés-Pujolràs et al., 2025; Twomey et al., 2016) sustain until response execution. Note that, in our simulations (*Fig. 5A,D*), the average traces do not always reach the estimated bound: this is because, in a fraction of trials, the bound was never hit, bringing averaged values away from the bound value.

**Figure 5.**
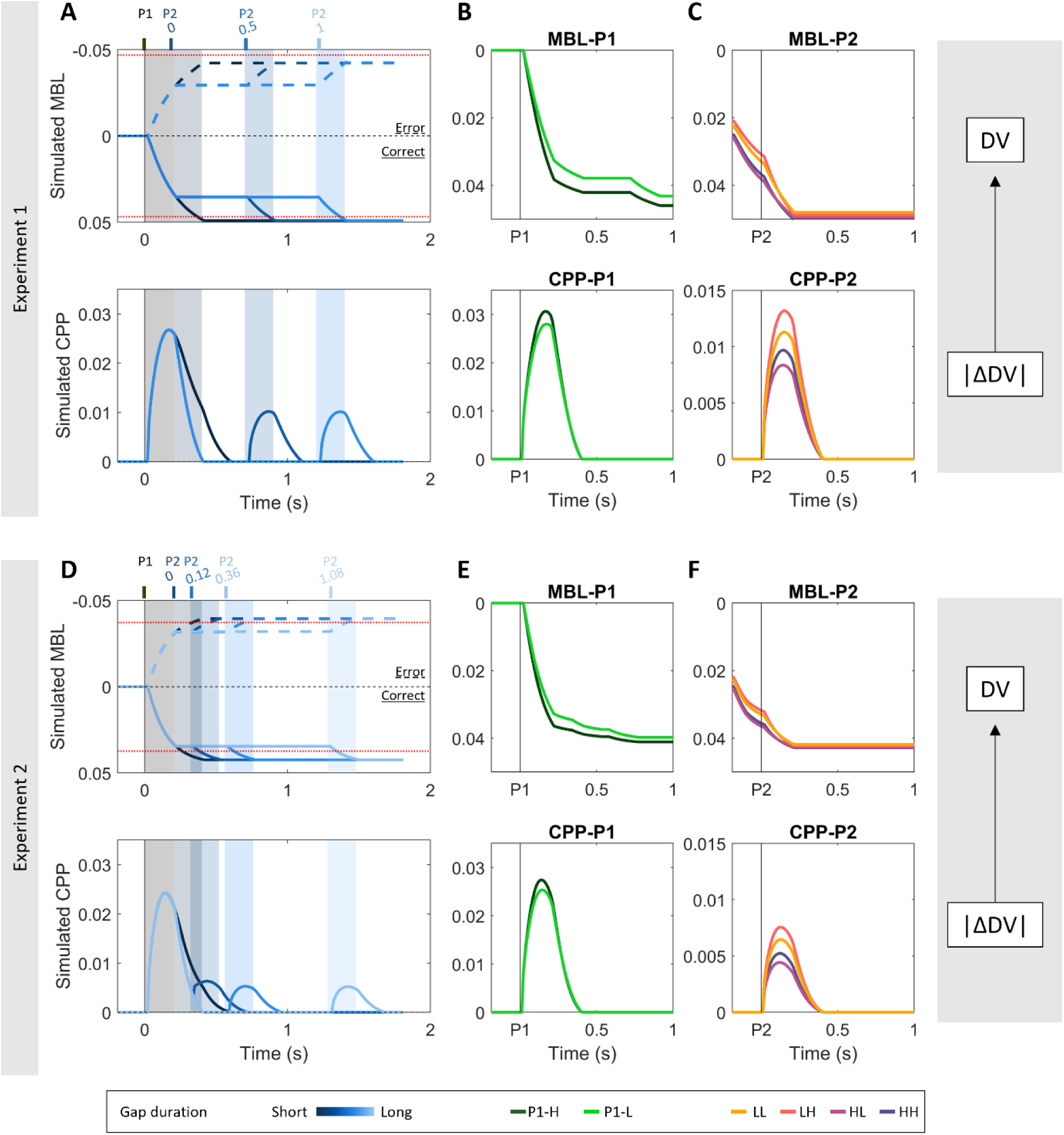
Model-derived simulations of motor beta lateralisation (MBL), reflecting the overall ‘DV’, and centroparietal positivity (CPP), reflecting local pulse accumulation (|ΔDV|) recapitulate empirical data. **A,D.** Simulated average MBL (*top*) and CPP (*bottom*) traces over the whole trial, sorted by gap duration. MBL traces are additionally sorted by accuracy (solid line = correct; dashed lines = error). The simulations recapitulate the bimodal shape of the CPP across the whole trial, with the second pulse reaching lower amplitudes than the 1st one in both experiments (see Fig. 2B). In turn, by assuming that MBL signals track the state of the DV throughout the trial and because no leak or decay is assumed, our simulations could also recapitulate the sustained nature of motor lateralisation signals throughout the gap. Note that DVs were simulated so that positive signs are associated with correct choices, and negative signs to errors (hence the y-axis inversion to accord with empirical MBL signals). Dotted red lines indicate the bounds for error/correct trials, respectively. **B,E.** Simulated MBL-P1 (top) and CPP-P1 (bottom), sorted by P1 coherence, and averaged across all gap durations and accuracies. Note that the step-like changes in MBL traces are due to averaging the DV values over the various gap durations. No such effect is observed in simulated CPP potentials because traces are derived from P1 activity exclusively. **C**,**F.** Simulated MBL-P2 (top) and CPP-P2 (bottom), sorted by P1 and P2 coherence, and averaged across all gap durations and accuracies. Our simulations recapitulate the observed coherence effects at both CPP-P1 (see Fig. 3A,C) and CPP-P2 (see *Fig. 3E,G*). Note that, for CPP simulations, we plot the estimated build-up at P2 assuming that activity started at 0 and not considering any potential overlap from post-P1 fall-down earlier on in the simulated trial, for consistency with the empirical traces where CPP-P1 activity is removed from the P2-evoked traces. In contrast, the MBL-P2 traces continue from their post-P1 state, accounting for the sustained nature of the signal (see *Fig. S2,S3* for empirical MBL-P1/P2 traces). Note also that, for the sake of simplicity, our CPP simulations do not assume any delay between stimulus presentation and signal build up. Therefore, our simulated CPPs start building up immediately after pulse onset, while empirical ones start with approximately a 0.2s delay. Similarly, although motor preparation signals lag the motor independent buildup dynamics at the CPP level, we assumed instant transmission here for simplicity. Since only accuracy and not reaction time was to be captured in the modelling of this delayed-response task, it was not necessary to estimate transmission delays.

In turn, our results suggested that the transient CPP activity reflected local, pulse-driven accumulation fed to the DV while evidence was present. This accumulation process stopped during incoherent random dot intervals, leading to the signal falling down rather than maintaining the global DV state. Thus, we simulated CPP activity by taking the absolute values of the cumulative evidence gathered within each pulse (|ΔDV|) on single trials (in keeping with the always-positive nature of the CPP; see Afacan-Seref et al., 2018; Kelly et al., 2021) and averaging across trials. If a bound was hit by the overall DV, the CPP traces were set to return linearly to 0 within 200 ms, as empirically observed in action-locked traces in self-paced tasks (e.g. Steinemann et al., 2018), and it did not resume upon presentation of a second pulse. This fall-down is only a signal feature accompanying the deactivation of the intermediary integration process and does not bear on choice. Given that the CPP traces in our grand-averaged data (*Fig. 2B*) fell fully back to baseline by the end of the longer gaps, CPP traces were also assumed to linearly return to zero as soon as the dots turned blue at the end of evidence pulses if a bound had not been hit by that time. Such a pattern assumes that the CPP generators are active during periods where evidence is presented (yellow dot pulses), but become inactive and fall down to baseline once evidence stops (blue dot periods). During pulse processing, the CPP feeds its output to the downstream motor level, which sustains its current level during the gap rather than deactivating. In this way, the motor process continuously represents the overall DV throughout the trial: the evidence that had been integrated by the CPP during pulse 1 (ΔDV_P1) is maintained at the motor level through the gap, is potentially updated by another round of accumulation by the CPP during pulse 2 provided the bound was not already reached, and thereafter further sustains up until response execution. Given the absence of previous empirical data on how signals may fall down without having hit a bound, we assumed the same fall-down rate in this case as when a bound is hit (linear return to zero within 200 ms).

The simulated single-pulse accumulation traces from our model captured the sensitivity of both CPP-P1 and CPP-P2 to motion coherence. The difference in CPP-P1 slopes for high vs. low coherence pulses is simply explained by the high vs. low drift-rate associated with these conditions (*Fig. 5B,E*). In the more complex case of CPP-P2, the activity evoked by P2 reached higher amplitudes in trials where P1 was low coherence (*Fig. 5C,F*). Our model can account for this due to two effects: First, because P1-Low trials are less likely to terminate early (*Fig. 4B*), and therefore fewer trials take a value of zero during P2. Second, because among those trials that do not terminate early, the level reached by the DV by the end of pulse one (ΔDV_P1) is on average lower for low-than high-coherence (*Fig 4C; Fig 5B,E*), and therefore more evidence from P2 needs to be integrated before the bound is hit and accumulation terminated. Thus, the higher CPP-P2 amplitudes reached in trials with low P1 coherence reflected both a higher proportion of trials where participants engaged in accumulation of P2, and the fact that they had to cover a longer distance to reach the bound (i.e. larger changes in the DV (|ΔDV_P2|) were possible before a bound was hit).

The model also predicted differences in CPP amplitude in correct/incorrect trials, which our dataset broadly recapitulated (*Fig. S6*). In particular, it predicted stronger CPP-P1 coherence-dependent effects in correct vs. incorrect trials, and it also predicted an interaction between accuracy and P2-coherence on CPP-P2 amplitude in trials with mixed coherences (HL, LH).

Finally, we assessed the extent to which some of our models’ assumptions influenced the simulated CPP profile. For example, we assumed for parsimony that the CPP would deactivate with the same temporal dynamics when a decision process terminates because a bound has been hit, as when accumulation is reset at the start of the gap, but we do not know this to be the case. In fact, our simulations show the CPP falling down rapidly to baseline in all gap conditions, while empirical data show the CPP reaching all the way to baseline for the longest gap condition only. We therefore conducted additional simulations to illustrate how different assumptions about the fall-down duration (i.e. time to go back to 0) and delay-to-fall (i.e. the amount of time elapsed before the signal stops accumulating and starts going down) would affect the CPP’s profile during the gap in our two experiments (*Fig. S7*). These parameters affected the CPP shape during the gap, but the patterns of coherence-dependent CPP-P1 and CPP-P2 amplitudes remained unchanged. Longer fall-down durations better recapitulate the observed empirical fall-down in short gap conditions, where signals fail to reach baseline before the second pulse onset. These simulations suggest that our assumption of a 200ms fall-down time may have underestimated that of the true signal dynamics, but dedicated experiments will be required to precisely quantify the fall-down characteristics of the CPP in these scenarios. In turn, allowing for some post-decisional evidence accumulation before the signal starts falling down also better captures some features of our empirical data, such as the higher amplitudes and longer peak latencies observed in the zero gap condition (*Fig S7*). We also simulated an alternative scenario where the signal is not assumed to fall (*Fig. S8*), which would be predicted if the CPP’s neural generators remained active throughout the gap, encoding a sustained representation of the DV, so long as the bound has not been crossed. These simulations did not recapitulate the CPP’s full drop to baseline in the long gap conditions, and produced distinctly smaller CPP-P2 evoked potentials relative to CPP-P1 (Exp. 1: CPP-P2 = 19% of CPP-P1 amplitude; Exp. 2: CPP-P2 = 13% of CPP-P1 amplitude) than observed in the empirical data (see *Fig. 2B*), which were more closely approximated by the model assuming a full reset during the gap (Exp. 1: CPP-P2 = 53% of CPP-P1 amplitude; Exp. 2: CPP-P2 = 35% of CPP-P1 amplitude; see *Methods* for full details on this analysis). Thus, the data are more consistent with a full reset of the intermediate accumulation process reflected in the CPP at the end of the pulse containing informative evidence, regardless of it having crossed a bound, with the maintenance of intermediate, potentially subthreshold levels of the overall DV being encoded solely at the downstream level of motor preparation.

Only one of the alternative flexible weighting models we considered could explain both the behavioural and neural patterns of data we observed - a model where the second pulse is downweighted as an inverse function of the strength of the first pulse. However, this model was not favoured by the model comparison approach (see Supplementary Note 1).

## 3. Discussion

A wealth of studies has investigated the neural mechanisms underpinning evidence accumulation in tasks where information is continuously available, identifying multiple neural decision signals at various levels in the sensorimotor hierarchy (Gold & Shadlen, 2007; Heekeren et al., 2008; O’Connell & Kelly, 2021; Okazawa & Kiani, 2023). However, little is known about the mechanisms underpinning decision-making in contexts where evidence from a given source comes in temporally separate bouts. This is important not only because many real-life decisions require this kind of integration, but also because neural decision signals can exhibit functional dissociations when probed in more complex contexts (Okazawa & Kiani, 2023; Parés-Pujolràs et al., 2025). In this study, we asked: how do neural decision signals support evidence accumulation if information availability is paused for a variable amount of time? Focussing on two human EEG decision signals, the centroparietal positivity (CPP) and motor beta lateralisation (MBL), we show that they reflect processes with distinct roles during decision formation in this context. Across two experiments, we show that the CPP does not exhibit sustained activity during the gap between evidence pulses, but exhibits transient build, peak and fall dynamics in response to each pulse that track distinct rounds of evidence accumulation, and reflect the absolute contribution of each pulse to the overall decision variable (|ΔDV|). In contrast, MBL exhibited build-up during pulses and sustained activity during the gap between, consistent with the encoding of a signed DV evolving across pulses. Our behavioural and neural data can be explained by a simple bounded accumulation process, where participants sometimes terminate decisions early despite a cost in accuracy.

Our finding that the CPP exhibits transient activations tracking multiple rounds of evidence accumulation that feed into a single decision provides further insight into its functional role. Most previous studies using continuous evidence paradigms showed that the CPP steadily grows throughout an evidence accumulation process, in parallel with motor preparation, consistent with its reflecting the absolute value of an underlying, continuously-accumulating DV (Kelly & O’Connell, 2013; O’Connell et al., 2012; Steinemann et al., 2018). In contrast, in our task the CPP exhibits two distinct build-up processes that align with the two pulses presented in each trial. This suggests that the CPP builds up while accumulation is ongoing, and disengages when the accumulation process is paused or a bound is hit. This interpretation of our data was further supported by simulations of the CPP profile under various signal decay assumptions, which showed that empirical signals were better recapitulated by a process that assumes that the CPP signal falls down toward baseline when evidence is paused (*Fig S8*). Furthermore, we observed that the amplitude of each pulse-evoked CPP reflected the weight of that pulse on choice. At the grand average level, the initial pulse systematically evoked higher amplitudes than the second one, consistent with its greater influence on choice behaviour (*Fig. 2B*). Further, single-trial regressions confirmed that fluctuations in CPP-P1 amplitude for a given coherence level were linked to behavioural variability, with higher-than-average CPPs associated with a stronger weighting of the first pulse on choice *(Fig S11).* Put together, these results are suggestive of a direct link between the CPP amplitude following a particular stimulus and that stimulus’ weight on choice. Empirical observations from previous studies align with this interpretation. In a previous study using the same gaps task (Azizi & Ebrahimpour, 2023), it was the second pulse that had a stronger weight on choice rather than the first (discussed below), and we note that CPP amplitudes were accordingly larger for pulse 2. Thus, although the direction of the behavioural bias differed across studies, in both, the bias in CPP amplitudes was in the same direction as the behavioural bias.

Our results support the idea that the CPPs following sequential pulses capture multiple rounds of evidence accumulation that are part of a single decision, and that its amplitude reflects each pulse’s partial contribution to an evolving DV. In contrast to the transient pattern of activity that characterises the CPP in our task, MBL started shortly after the first pulse, and was sustained at a coherence dependent level throughout the whole trial duration (*Fig 2 & Fig S2*). This is in line with multiple previous studies, which have identified sustained encoding of decision variables in motor preparatory signals (Donner et al., 2009; Murphy et al., 2016, 2021; Parés-Pujolràs et al., 2025; Wyart et al., 2015). Put together, our findings support the idea that in this task the CPP feeds to accruing evidence from each individual glimpse of evidence (|ΔDV|) to downstream motor signals that maintain a sustained representation of the evolving DV. Thus, our study identifies new task-dependent functional dissociations between distinct levels of the neural architecture underpinning decision making.

The dynamics within these levels and interactions between them were further illuminated by our computational models. In our two experiments, both the behavioural and neural effects could be reproduced with a two-level bounded accumulation model, in which an intermediate accumulator operates locally on contiguous periods of evidence, feeding its output on to a thresholded motor level where the cumulative total is sustained. Our model assumed that participants stop accumulating evidence if a bound is hit and that accumulation is paused during the gap between pulses; while the cumulative total is held at the motor level, the intermediate accumulator resets. By assuming that the CPP becomes inactive and falls down during the gaps, our simulations could recapitulate key CPP characteristics observed in our task including the opposite effects of the first and second pulse coherence on its amplitude. The fact that trials with low coherence first pulses were less likely to be terminated early, and had to cover a longer distance-to-bound upon presentation of the second pulse explained why CPP-P2 reached higher amplitudes in those trials.

A key feature of our model which enabled it to account for our behavioural observations was that accumulation was bounded, so that decisions could be terminated before all of the evidence has been sampled and accumulated. Given that the motion in any two sequential pulses in the gaps task always went in the same direction, that behavioural responses were deferred until a later cue, and that single-pulse accuracies were far from ceiling, the optimal strategy would have been to perfectly accumulate all available evidence; by setting a bound, our participants often missed some information and thus achieved suboptimal accuracy. While behaviour approximating perfect integration in relatively long timescales has been reported across species (Brunton et al., 2013; Waskom & Kiani, 2018), suboptimal integration has also been frequently observed. Previous expanded judgement tasks prescribing perfect integration have often reported recency biases (Cheadle et al., 2014; Wyart et al., 2015), but primacy effects consistent with early decision terminations have also been observed in both humans (Balsdon et al., 2020) and non-human primates (Kiani et al., 2008; Mazurek et al., 2003; Yates et al., 2017). Some studies have even found “bump-shaped” psychophysical kernels indicating increased weighting of evidence occurring around the middle of a stimulus sequence (Keung et al., 2019, 2020; Sospedra et al., 2025). Such deviations from optimal integration may reduce cognitive effort at the cost of accuracy, and may emerge due to multiple factors, including stimulus properties (Bronfman et al., 2016; Lange et al., 2021), biases driven by stimulus consistency (Cheadle et al., 2014; Glickman et al., 2022) or choice history (Urai et al., 2019), or result from different integration strategies that may be flexibly deployed, such as leaky (Tsetsos et al., 2012) or bounded accumulation (Kiani et al., 2008; Mazurek et al., 2003).

In the context of our task, previous studies using the same stimuli found the opposite bias in behavioural weighting, where the second pulse had a stronger influence on choice than the first (Azizi & Ebrahimpour, 2023; Kiani et al., 2013). Given that the stimulus characteristics (dot speed, density) in previous work were the same as in our study, differences in trial structure, difficulties and training may account for this discrepancy instead. For example, our task had comparatively long lead-in periods before the first pulse (min. 0.7s compared to min. 0.4s of previous studies), which may have helped them to prepare for full integration of the first pulse. In addition, while previous studies included at least four coherence levels, we used only 2, which possibly lends itself to calibrating a bound to achieve acceptable accuracy while saving cognitive effort. Furthermore, we did not use trial-by-trial feedback but rather provided aggregate performance metrics at the end of each block, which would have detracted from participants’ ability to monitor their own performance. Finally, even though the probability of a single pulse trial was the same, the single-pulse staircasing procedure we used at the beginning of the task may have encouraged a strategy more focused on the information in the first pulse. In line with this interpretation, a recent study that also lacked trial by trial feedback and used a single-pulse training procedure has also reported recency effects in a task with different visual stimuli but the same variable-delay task structure (Golmohamadian et al., 2025). However, resolving the factors that encourage early termination strategies will ultimately require dedicated experimental manipulations.

Our findings regarding the CPP’s contribution to decisions based on temporally separate bouts of evidence aligns with the results of recent work in expanded judgement tasks (Parés-Pujolràs et al., 2025; Wyart et al., 2012, 2015). In expanded judgement studies, participants view a sequence of discrete, easily-identified tokens that support one of two choices and must integrate over multiple tokens to make an accurate decision. In these tasks, CPP responses to discrete tokens have been shown to scale with context-optimal belief updates in classic non-volatile contexts that prescribe perfect linear accumulation (Wyart et al., 2012, 2015) as well as in volatile environments where the correct answer may change within a given trial, which prescribe a nonlinear transformation of the token information being integrated (Parés-Pujolràs et al., 2025). That is, evoked CPP activities track how much a DV changed following each piece of evidence, but not the absolute running sum of accumulated evidence. This stands in contrast to the results from conventional decision-making tasks where evidence is presented continuously, and in the form of high-noise continuous stimuli (Geuzebroek et al., 2023; Gherman et al., 2024; O’Connell et al., 2012; Steinemann et al., 2018) rather than low-noise discrete tokens. Our task represents a middle ground between these two types of paradigms: we used noisy continuous evidence, but grouped it in two temporally discrete, sequential windows. Thus, participants needed to integrate over time to discriminate the information in each evidence pulse, but also across evidence pulses to maximise their accuracy. This allowed us to show that, in conditions where evidence is noisy and continuous but can be paused, the CPP’s profile separately tracks multiple rounds of evidence accumulation. When such updates occur in discrete steps as in expanded judgement tasks, or in short dynamically-accruing bouts as in our gaps task, the CPP does not seem to keep track of the state of an overall DV between temporally separated pieces of evidence, but rather processes each piece separately. However, it remains unclear to what extent the CPP may behave differently in other contexts. In particular, a common feature between expanded judgement tasks and our design is that the presence of evidence is unambiguously indicated, in our case by colour cues. These colour cues may have acted to trigger initiation and termination of two rounds of evidence accumulation. However, it is plausible that in the absence of such external cues the CPP may have exhibited a sustained pattern of activity consistent with continuous monitoring of the environment (Geuzebroek et al., 2023), potentially plateauing during periods with incoherent dot motion that should elicit close-to-zero, or heavily reduced, momentary belief updates on average. This will require a dedicated experiment to test.

Our study also leaves some open questions. Given that the coherence-dependent effects of gap duration on behaviour were not expected and were inconsistent across the two experiments (*Fig 1B*), we did not aim to capture them in our modelling. However, surmising that they might relate to dynamic expectations arising from the temporal structure of the task, we sought to gain insight into dynamics of attentional engagement and expectation through exploratory analyses of occipital alpha activity (*Fig 2C, Fig S9*). Occipital Alpha-band power has been linked to attention and temporal expectations (Rohenkohl & Nobre, 2011), and also to the onset (Van Den Brink et al., 2021) and build-up rate (Kelly & O’Connell, 2013) of evidence accumulation signals like the CPP. In our two experiments, the gap durations between the two pulses followed different distributions: in Exp 1 gaps were uniformly distributed, whereas in Exp 2 they were skewed towards shorter gaps durations. We found evidence that occipital alpha power was sensitive to this difference (*Fig S9*), in that it was relatively reduced during the longer gaps in Exp 1, where they were relatively more likely, compared to Exp 2. We also found that phasic alpha decreases additionally modulated the weight of the second pulse on behaviour in Exp 2 (*Fig S11*), suggesting that phasic alpha activations may influence evidence integration. However, more work is required to specifically test how tonic and phasic occipital Alpha may interact with evidence accumulation processes, or influence behaviour through other mechanisms.

While the bounded model provides an intuitive account of the push-pull relationship whereby the more evidence is used from P1, the less is used from P2, it was important to explore alternative models based on the idea that the evidence itself is unequally weighted across the two pulses. Previous studies where sequential samples may either be consistent or inconsistent with an evolving decision variable (Cheadle et al., 2014; Glickman et al., 2022; Park et al., 2025) have reported consistency biases, where evidence supporting current beliefs is boosted rather than dampened. In all these studies, the opposite alternative to that which gained early support is selectively downweighted. However, in our task the two pulses always favoured the same alternative, so that no inconsistent evidence was ever presented. Thus, it is plausible that participants may have partially withdrawn attention from the task following the first pulse, and especially so if evidence was strong. Based on this idea, we devised an unbounded model in which P2 drift rate is downscaled as a function of the coherence of P1. This could account for both behavioural and neural results, producing a primacy effect and a pattern of CPP-P2 amplitudes that recapitulated the observed pattern (see Supplementary Note 1). While this model was not supported by quantitative model comparison (Fig. S15), and it does not change the key finding that CPP-P2 amplitude mirrored its contribution to choice, the fact that it could at least capture the key patterns qualitatively calls for discussion of its plausibility. In the bounded model, the integration of the second pulse is curtailed or entirely precluded as a function of the evidence accrued by the end of P1 and hence the remaining distance to the decision-terminating bound, if any. These early terminations before or during P2 thus save any energy associated with further integration, at a cost to accuracy. In contrast, the downweighting model entails fully integrating P2, but in a dampened manner with reduced SNR.

In sum, our study provides novel insight into the neurocomputational mechanisms supporting decision making in a context where decisions need to be made on the basis of a single stimulus feature that is only intermittently available. Our task allowed us to identify novel functional characteristics of a well-established neural decision signal in the human EEG: the CPP. We show that in tasks where perceptually noisy evidence is presented intermittently in short bouts, the CPP does not maintain a sustained representation of cumulative evidence, but rather engages in multiple rounds of evidence accumulation during those bouts, which in turn feed a continuously updating, sustained DV represented in downstream areas. This local evidence-processing property is consistent with recent work showing that the CPP encodes belief updates rather than the ongoing state of a DV in expanded judgment tasks involving the integration of a series of stimuli that are individually easy to identify (Parés-Pujolràs et al., 2025), and suggests that the CPP may only trace the entire evidence accumulation process underpinning decisions when the physical evidence is presented in one continuous stream (Geuzebroek et al., 2023; Kelly & O’Connell, 2013; O’Connell et al., 2012; Steinemann et al., 2018). Our results thus add to growing work suggesting that the neural architecture underlying evidence accumulation for decision-making can exhibit flexible functional roles in different contexts, underpinning adaptive behaviour.

## 4. Methods

### Participants

Twenty-three participants were recruited to take part in Experiment 1, and 23 different participants were recruited to take part in Experiment 2 of this study. In Experiment 1, one participant withdrew after a few blocks of the task because they were feeling dizzy, and thus their data were excluded from further analysis. Two participants were unable to perform the task at the minimum required level of accuracy and were excluded from analysis in Experiment 2. A total of 22 (M = 21.5 years old, SD = 1.79 years old, 10 females) and 21 (M = 22 years old, SD = 3.85 years old, 11 females) participants were thus included in the final samples for Experiment 1 and 2 respectively. All participants provided informed written consent before participation in the study and were compensated for their time (10€/hour), and all procedures were approved by the University College Dublin Human Research Ethics Committee.

### Task

Participants performed a motion direction discrimination task adapted from Kiani et al. 2013 using random dot kinematograms (RDK). The experiments consisted of a total of 700 (Exp 1) and 933 trials (Exp 2), divided into blocks of 70 trials each. Most trials in the experiment were two-pulse trials (85.7%) and the remaining fraction were single-pulse trials (14.3%). In single-pulse trials, the second pulse was not presented. Instead, the first pulse was followed by a 0% coherence lead-out period and then the trial terminated. The proportion of single- and two-pulse trials mirror the ones in the original Kiani et al. 2013 study.

Stimuli were presented centrally on a standard monitor, and participants sat at a 50-60cm distance from the monitor. Dots were presented within a 5° circular aperture at the center of the screen, the dot density was 16.7 dots/degree^2^/s, and coherent dots moved at a speed of 5 degrees/s. Due to a technical error, the monitor refresh rate was 60Hz for 19 participants and 85Hz for 3 participants in Experiment 1, while it was 85Hz for 19 participants and 60Hz for 2 participants in Experiment 2. Each trial began with a white fixation cross presented at the centre of the display on black background. Then, a patch of 0% coherence dots appeared in blue colour on the centre of the screen (700 – 1050 ms, with short (700/750ms), medium (900/950ms) or long (1000/1050ms) delays appearing with a probability of 0.7,0.2 and 0.1, respectively). This was followed by a 200 ms pulse of yellow dots moving at either high or low coherence. After a gap of variable duration (Exp 1: [0, 0.5, 1] s, Exp 2: [0, 0.12, 0.36, 1.08] s) a second 200 ms pulse of either high or low coherence motion was presented. Finally, a patch of blue dots moving at 0% coherence was presented (700 to 1050 ms) in blue. The fixation cross was visible throughout the trial to encourage participants not to follow the dots with their eyes. At the end of the final blue dot sequence, all stimuli disappeared. Trials were separated by a fixed 1s intertrial interval. All gap durations were equally likely, and all possible motion coherence combinations (low-low, low-high, high-low, high-high) and gap durations were randomly permuted throughout the experiment. Three gap durations were originally used in Experiment 1 in order to maximise trial count for each of a small number of gap durations. Having observed that the participants utilised the second pulse less than the first, opposite to previous work (Kiani et al., 2013), we ran a second experiment with shorter gap durations made more prevalent, on the basis that closer temporal contiguity might make it less likely that participants would disengage from accumulation before the second pulse was presented and encourage continued accumulation. The expectations for a long gap to occur differed in our two experiments. Given the percentage of 1-pulse trials, if the second pulse had not yet occurred 500 ms after the 1st pulse the probability of it happening was higher in Experiment 1 (p = 0.666 [1/3*85.7/(1/3*85.7+14.3)]) than in Experiment 2 (p = 0.6 [1/4*85.7/(1/4*85.7+14.3)]. Participants were told they had to respond after stimulus offset, with no time limit. Participants responded by pressing the “C” or “M” keys on a standard keyboard with the left and right thumb respectively. The blue and yellow colours used in the experiment were selected to be physically isoluminant, as measured on the lab monitor, in order to avoid large evoked potentials arising from strong luminance changes.

### Staircase procedure

In order to estimate the required dot coherence for participants to perform at 70% accuracy across coherence levels, all participants performed a staircase procedure consisting of 100 trials. All trials consisted of 0% coherence lead-in dots (700 to 1050 ms), a single pulse of evidence (200 ms) and a final sequence of 0% coherence lead-out dots (700 to 1050 ms). The staircase followed a 2-down, 1-up procedure (Levitt, 1971), and the starting value was set to 40% for all participants. After two correct responses, the coherence was lowered. The initial step size was set to 6%, and it was halved every second time the staircase went down. This was done to achieve greater precision towards the end of the staircase. The average coherence of the dot motion in the last 40 trials of the staircase procedure were used to estimate the two coherence strengths used in the task. Specifically, high (Exp. 1: M = 44.73%, SD = 12.64%; Exp 2: M = 33.04%, SD = 13.41%) and low (Exp. 1: M = 26.75%, SD = 7.74%; Exp 2: M =19.82%, SD = 8.04%) coherences were then set to be 25% higher or lower than the staircase-estimated coherence.

To keep performance constant throughout the task, the average accuracy was checked after each block and coherence was dynamically adjusted between blocks. If accuracy was on average lower than 60% or higher than 90% across conditions, coherence was adjusted by increasing or decreasing it by a tenth of the starting value obtained from the staircase procedure. This precaution was taken to avoid ceiling & floor effects. Most participants’ performance was stable throughout the task, and only a few required substantial adjustments of coherence between blocks (see *Fig. S1* for individual data).

### EEG recording and preprocessing

Continuous EEG was recorded from 128 scalp electrodes using a BioSemi system with a sample rate of 512 Hz. Eye movements were recorded with four additional electro-oculogram (EOG) electrodes, two above and below the left eye and two at the outer canthi of each for vertical (VEOG) and horizontal HEOG) eye. Additionally, we recorded electromyogram (EMG) signals from the thenar eminence using two electrodes on the left and two on the right hand, as a standard procedure, though EMG was not of interest in the current study. All data were analysed in a Matlab R2021a using the EEGlab (Brunner et al., 2013; Delorme & Makeig, 2004) and the ERPlab plugin (Lopez-Calderon & Luck, 2014).

Continuous data were high-pass filtered at 0.1Hz, and low-pass filtered at 40Hz using a zero-phase shift Butterworth filter. Then, stimulus-locked epochs were generated from -1.5s to 3s around the onset of the first evidence pulse. Bad channels were defined as those exceeding 50 standard deviation units, and were interpolated. Data were then re-referenced to the common average of all EEG channels. Trials where participants blinked during one of the evidence pulses were excluded from analysis. Eye movements occurring at any other time during the trial were removed from the data using Independent Component Analysis (ICA). Epochs were baseline-corrected relative to the 200 ms before the onset of the first evidence pulse, and channels showing ±150µV deviations were interpolated on a trial-by-trial basis using the TBT toolbox (Ben-Shachar, 2018). If any given epoch had more than 20 bad channels, it was marked for rejection. Further, if any given channel was marked as “bad” on >30% of epochs, it was interpolated across the whole experiment. Finally, we applied a Current Source Density (CSD) transform (second spatial derivative) using the CSD toolbox (Kayser & Tenke, 2006) to the preprocessed data to reduce the spatial spread and thereby reduce overlap between topographic foci. Time frequency decomposition was performed using a 7-cycle morlet wavelet transform, extracted in 1-Hz steps in the 1-40Hz range. Alpha and beta band power were obtained by averaging power in the 8-12Hz and 13-30Hz frequency ranges, respectively.

### Behavioural analysis

First, to investigate the effect of pulse coherence and gap duration on participants’ accuracy, we ran the following logistic regressions on one-pulse (*Eq. 1*) and two-pulse trials (*Eq. 2*):

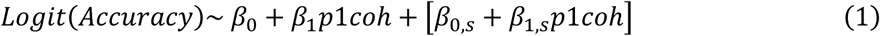

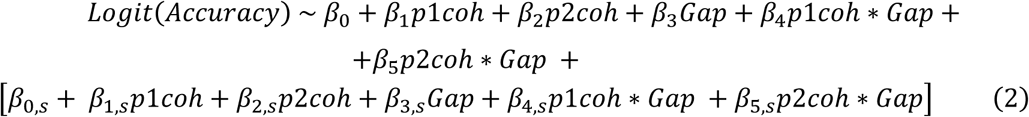

Where terms between square brackets indicate random effects; *s* refers to each participant, and indicates that random effects (intercepts, or slopes) were calculated in addition to fixed effects for that particular coefficient. Next, to test whether participants were generally more accurate in two-pulse compared to one-pulse trials we ran the following logistic regression (*Eq. 3*):

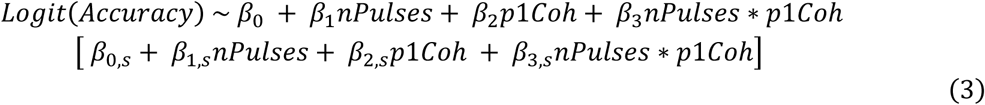

This analysis was restricted to two-pulse trials with equal coherences in the two pulses (HH, LL trials only) so that any effect of pulse number would be exclusively related to the presence of additional motion, rather than potentially capturing order or coherence-consistency related effects.

Then, to investigate the effect of pulse order on participants’ accuracy, we ran the following regression (*Eq. 4*):

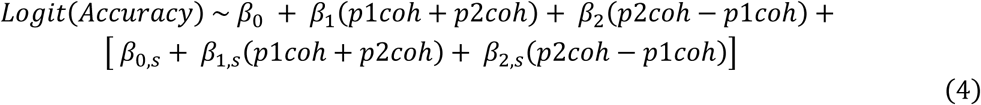

where the difference coefficient is expected to not be significant if the order of pulses does not influence performance.

In all regressions, coherence values were normalised within participant because they were staircased for each participant individually and varied widely. All models included a maximal random effects structure, with a random intercept and random slopes for all fixed effects to control for interindividual variability. All models were fit using the ‘bobyqa’ optimiser in the lme4 toolbox (Bates, 2011). In Experiment 2, this resulted in singular fits for both Eq. 2 and 3. Following previous suggestions, we used a weakly informative prior to regularise the random effects covariance matrix (Chung et al., 2015) using the blme package (Chung et al., 2013), which resolved the singular fit problem. In cases where the model output flagged potential convergence problems, we assessed the robustness of parameter estimates by comparing them across 6 different optimisers using the ‘allFit’ function in lme4 toolbox (Bates, 2011), as advised by the developers. All such model fits were deemed to be robust as the optimiser estimates rarely deviated by more than one decimal point from the ‘bobyqa’ ones.

Finally, we compared participants’ accuracy on two-pulse trials to the expected performance assuming perfect integration, and based on their accuracy in one-pulse trials. Following Kiani et al. 2013, we assumed that decisions are determined by the sign of the sum of evidence in the two pulses. The evidence distribution was inferred from participants’ accuracy on single-pulse trials as follows:

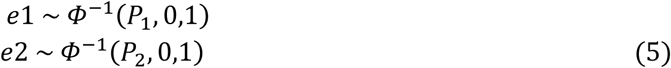

Where P1 and P2 are grand-average accuracies for high- and low-coherence one-pulse trials. **Φ** is the normal cumulative distribution function:

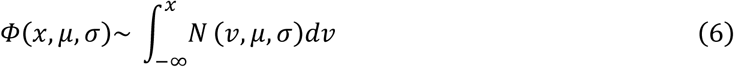

and **Φ**^-1^ is its inverse. N(v,μ,σ) is the normal probability density function with mean (μ = 0) and standard deviation (σ = 1). The expected accuracy for double-pulse trials was then calculated as follows:

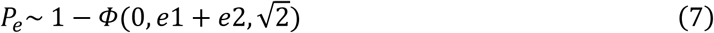

### ERP analysis

#### CPP-P1 & CPP-P2 analysis

To investigate how centroparietal responses to the two sequential pulses varied with motion coherence, we extracted EEG activity from a cluster of centroparietal electrodes where activity at 500 ms after pulse 1 was maximal across conditions (see *Fig 2*). To compute CPP-P1 slopes, we fit a line from 0.2s post P1 onset when the CPP appeared to begin building, to peak time for each participant individually and for each P1 coherence condition separately, restricting the peak search times around [0.4-0.6s] post-P1. We then compared the slopes in those two conditions by running a paired-samples t-test. For CPP-P2 visualisation, we removed P1-related activity by subtracting the traces from the longest gap conditions [-0.2 to 1.4s with respect to P1], from all traces. This resulted in an average EEG activity of 0 (see *Fig. S4*) preceding the second pulse, in all conditions, and rendered the key CPP-P2 findings more clearly visible by removing P1-induced EEG drifts. Note that this does not affect the CPP-P2 activity in the long gap conditions, as the subtraction alters data up to the end of P2 but not further (see *Fig. S4*). We report the average across all non-zero gap conditions in the main manuscript (*Fig. 2*), as well as the data sorted by gap in the supplementary material (*Fig. S3, S4*). We also used the so-corrected activity to compare the relative amplitudes of CPP-P1 and CPP-P2 across all non-zero gap conditions. For this, we first computed the average amplitude 0.4 to 0.6s after P1 (in the raw data, *CPP-P1_amp_*) and P2 (in the corrected data, *CPP-P2_amp_*), for each gap separately, and averaged across all non-zero gap conditions. Then, we computed the percentage of CPP-P1 amplitude reached by CPP-P2 as CPP-P2_amp_/CPP-P1_amp_ * 100.

To investigate the impact of dot motion coherence on both CPP-P1 and CPP-P2 amplitudes (*Fig. 3*), we ran a series of time-resolved regressions:

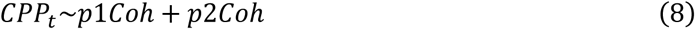

Where *t* indicates time from P1 for CPP-P1 analysis, and time after P2 for CPP-P2 analysis. For this analysis, we aligned all gap conditions to P1/P2 onset respectively and ran the regressions for all trials pooled, for each subject separately. For CPP-P2, data were additionally baselined using 100ms before P2 onset to minimise the impact of gap duration on average amplitudes. All EEG data were low-pass filtered (6Hz) prior to running the regression to reduce the impact of high frequency noise on these single-trial analyses.

Next, to investigate how trial-to-trial variability in attention and evidence accumulation influenced performance, we ran a series of time-resolved logistic regressions.

First, we investigated whether single-trial variability in centroparietal responses modulated the weight of pulses on choice. The interaction terms between pulse coherence and CPP amplitude in the following regression (*Eq. 9*) thus test whether pulses which evoked a particularly high centroparietal response were also associated with that pulse having a greater weight on choice:

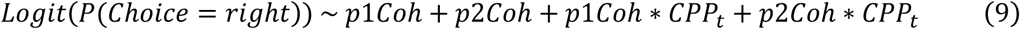

Where *t* indicates time after P1 for CPP-P1 analysis, and time after P2 for CPP-P2 analysis.

Second, we ran an analogous analysis to investigate whether fluctuations in evoked occipital alpha desynchronisation also modulated the weight of each pulse on choice (*Eq. 10*):

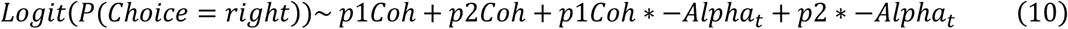

Where *t* indicates time after P1 for alpha-P1 analysis, and time after P2 for alpha-P2 analysis.

All regressors and EEG signals were z-scored across trials by timepoint. For alpha power analysis, we took the *negative* alpha power across a range of bilateral occipital electrodes (see highlighted electrodes in *Fig. 2*), so that positive coefficient values indicate that higher attention (i.e. lower alpha power) increase weight on choice or CPP amplitude. For the CPP analysis, we used the signal averaged over a cluster of centroparietal electrodes for CPP (*Fig. 2B*). Regressions were run for each gap condition separately for times between [0-1s] after each pulse, and the resulting coefficients were aligned to P1 and P2 and averaged across conditions.

#### EEG statistics

EEG evoked potentials and regression results were statistically evaluated against zero using cluster-based permutations using the FieldTrip toolbox (Maris & Oostenveld, 2007; Oostenveld et al., 2011). All tests were two-tailed, cluster-corrected (p<0.05), and 10000 permutations were run. We additionally compared the magnitude of the occipital alpha level in the longest gap across the two experiments using an independent samples t-test in Matlab.

#### Model fitting & simulations

In order to account for participants’ behaviour, we fit a simple diffusion model to the grand-averaged accuracy in the 4 conditions with three free parameters: bound (*b*) and two drift rates (*d1, d2*) corresponding to the low and high coherence conditions. The accumulator model followed this basic equation:

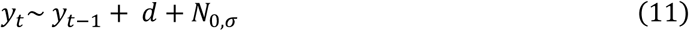

Where y_t_ is the accumulator output or decision variable (DV) for any time t within a pulse (where *d*=*d1* or *d = d2* depending on coherence). Gaussian noise, N_0,σ_, was added in order to represent noisy sensory evidence through the trial. During the gap, the accumulator was set to stay stable at its final within-pulse value so that *y*_t_ was set to equal *y*_t-1_ for any time *t* outside of pulses during which incoherent motion was presented and dots were blue. Given the behavioural observation that overall accuracy did not decrease in longer gap durations, and the fact that evidence pulses were clearly indicated by a change in the dots colour, the DV was paused and not allowed to further accumulate the noise during the 0% coherence motion gap separating the two pulses. When the DV crossed a bound ‘*b’,* evidence accumulation terminated. The accumulator assumed no noise or leak during the gap, and in any trials where the bound was not crossed accumulation continued upon presentation of the next pulse. Note that this one-dimensional, continuous, bounded accumulator represents the overall DV corresponding to the level of motor preparation. As we explain below, the CPP was simulated by extracting the single-pulse components of this model.

We fit the model by minimising G^2^ with the *particleswarm* function in the Global Optimisation Toolbox for Matlab. G^2^ is a goodness of fit measure that in this study quantified how much predicted accuracies differed from observed ones in each condition.

Computational models were fit to the two experiments separately simulating 10000 trials for each condition, and the best fitting parameters were then used to simulate CPP traces. Akaike Information Criterion (AIC) and Bayes Information Criterion (BIC) were computed to assess the goodness of fit as follows:

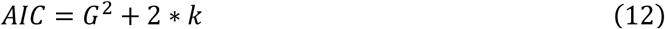

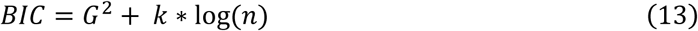

Where *k* indicates the number of free parameters for a given model, and *n* represents the number of trials used to fit the model.

### Neural data simulation

We simulated MBL activity by taking the evolving value of the simulated decision variable throughout the trial. If a bound was hit, the DV was set to stay stable in order to account for the sustained nature of MBL signals our empirical data as well as previous work in delayed response tasks (Murphy et al., 2021; Parés-Pujolràs et al., 2025; Twomey et al., 2016).

In turn, following previous work, we simulated the CPP by taking the absolute value of the underlying, signed accumulator traces unfolding during each individual pulse, produced by the best fitting parameters of our model (Afacan-Seref et al., 2018; Kelly et al., 2021). Taking the absolute value implicitly assumes that the differential evidence accumulator is implemented via two separate neural populations that encode values respectively favouring one or the other alternative. If the simulated decision variable hit a bound, the accumulator traces were set to fall back down to zero within 200 ms after hitting the bound, based on past observations of post-peak CPP dynamics (Steinemann et al., 2018). In the model shown in the main text, if the overall DV did not hit a bound by the end of the first pulse, it was assumed that the cumulative evidence level attained thus far was sustained in the overall DV at the motor level, but that the intermediate accumulator fell back to its zero baseline level, and restarted accumulation upon the presentation of the second pulse, which at the motor level added to the already sustained level. Variants of this scheme were also explored, as explained in *Fig. S7, S8*. In order to qualitatively compare how well various decay parameters matched the empirical CPP dynamics, we computed the relative CPP-P2/CPP-P1 percentage amplitude predicted by various models by extracting the average evoked amplitude of the simulated CPP [0-0.2s] for each pulse separately, with respect to the pre-pulse baseline amplitude. This 0.2s interval is analogous to the time used for analysis of the empirical data (see above), but is set to an earlier time period because our simulations did not incorporate any sensory delay, and therefore ramping activity started instantly after stimulus onset.

## Acknowledgements

This work was supported by a grant from Science Foundation Ireland (now Taighde Éireann) under grant number 19/US/3599, an Investigator Award from the Wellcome Trust under grant number 219572/Z/19/Z (to S.P.K) and a grant from the European Research Council (ERC): Consolidator Grant ‘IndDecision’ under grant number 865474. The authors would also like to thank Mike Shadlen for discussions on this work, and students Liang Tong and Bowen Duan for assistance during data collection.

## Author contributions

Conceptualisation: E.P-P, & S.P.K; software: E.P-P; data curation & analysis: E.P-P.; resources: S.P.K.; writing of original draft: E.P-P.; review & editing: E.P-P, R.G.O., A.C.G, & S.P.K.; supervision: S.P.K.

## Data & code

All raw EEG and behavioural data in this study as well as analysis code are publicly available in an OSF repository: https://osf.io/z7dp2/

## Supplementary material

**Figure S1.**
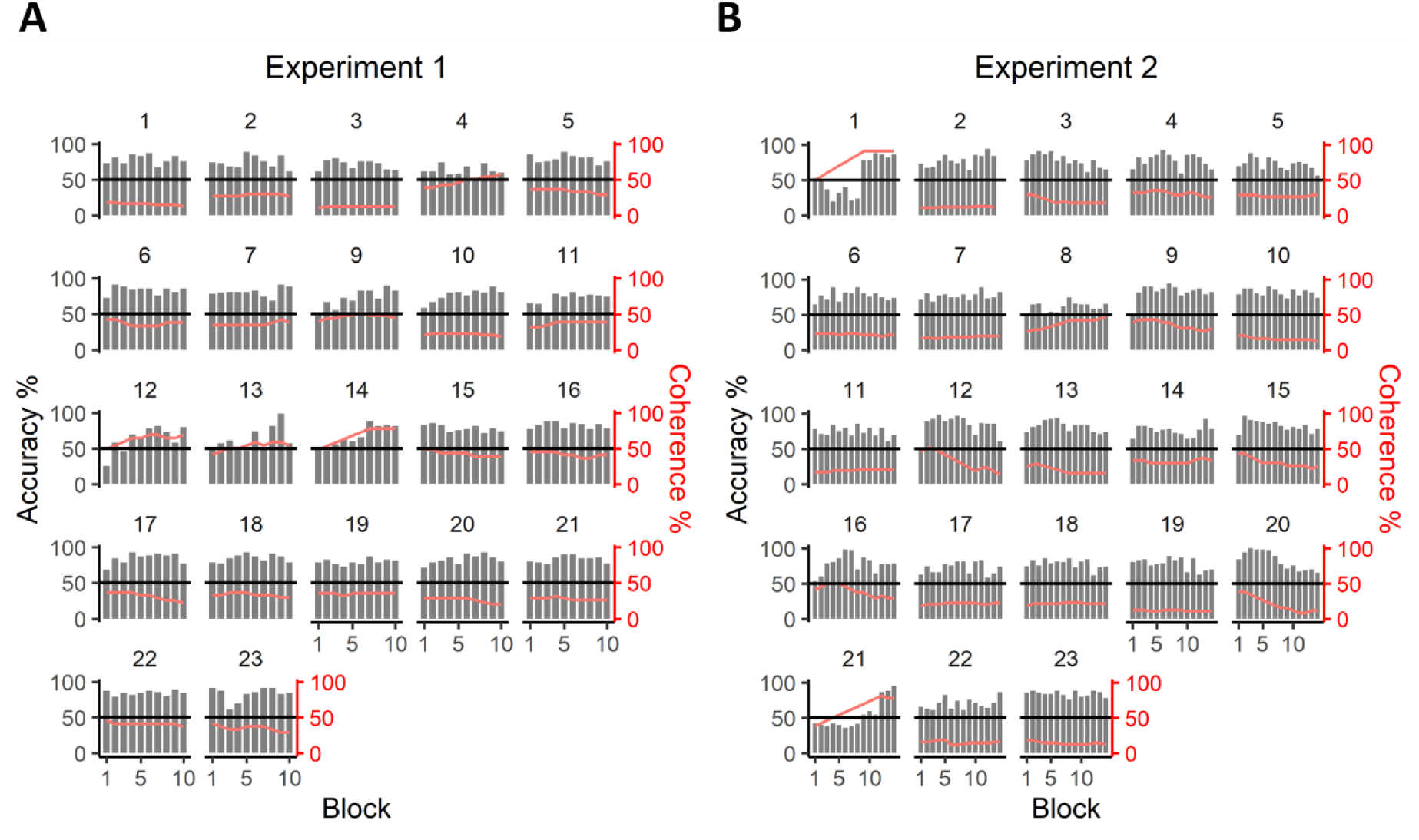
Individual participants’ accuracy (*grey bars*) and average coherence (*red line*) for each block in Experiments 1 (**A**) and 2 (**B**). On average, most participants performed close to 70% accuracy and clearly above chance (*solid black line*), and only required minimal coherence adjustments between blocks. The data of participants 1 and 21 in Experiment 2 were excluded from further analysis due to poor task performance in a majority of blocks.

**Figure S2.**
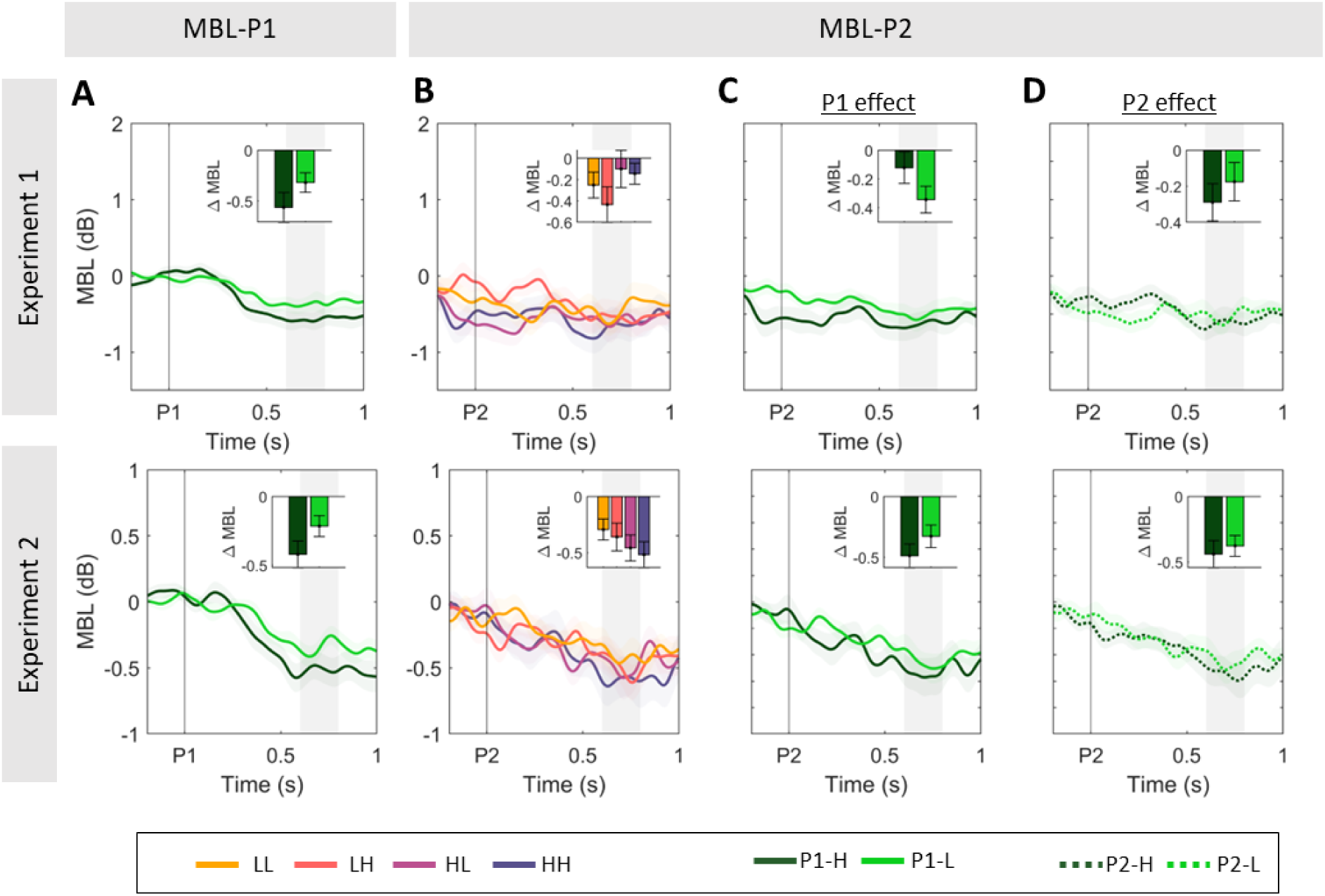
Sensitivity to evidence strength in the two pulses in motor beta lateralisation signals. **A.** Grand-averaged (±SEM) motor beta lateralisation [Contra-Ipsilateral hemispheres with respect to correct response hand] after pulse 1 (MBL-P1), averaged across all gap conditions and across all trials regardless of accuracy. Motor preparation started shortly after the first pulse, and was sustained at a coherence-dependent level. **B.** Grand-averaged MBL after pulse 2 (MBL-P2), averaged over all non-zero gaps in the two experiments. **C.** Effect of P1 coherence on MBL after P2. Traces illustrate the grand-averaged (±SEM) aligned to pulse 2 (MBL-P2), averaged over all non-zero gaps and across P2 coherence. **D.** Effect of P2 coherence on MBL after P2. Traces illustrate the grand-averaged (±SEM) aligned to pulse 2 (MBL-P2), averaged over all non-zero gaps and across P1 coherence. In panels **A-D,** inset bar graphs illustrate the MBL excursion as the difference between activity at [0.6-0.8s, shaded area] after pulse onset, minus activity at baseline [-0.2-0s].

**Figure S3.**
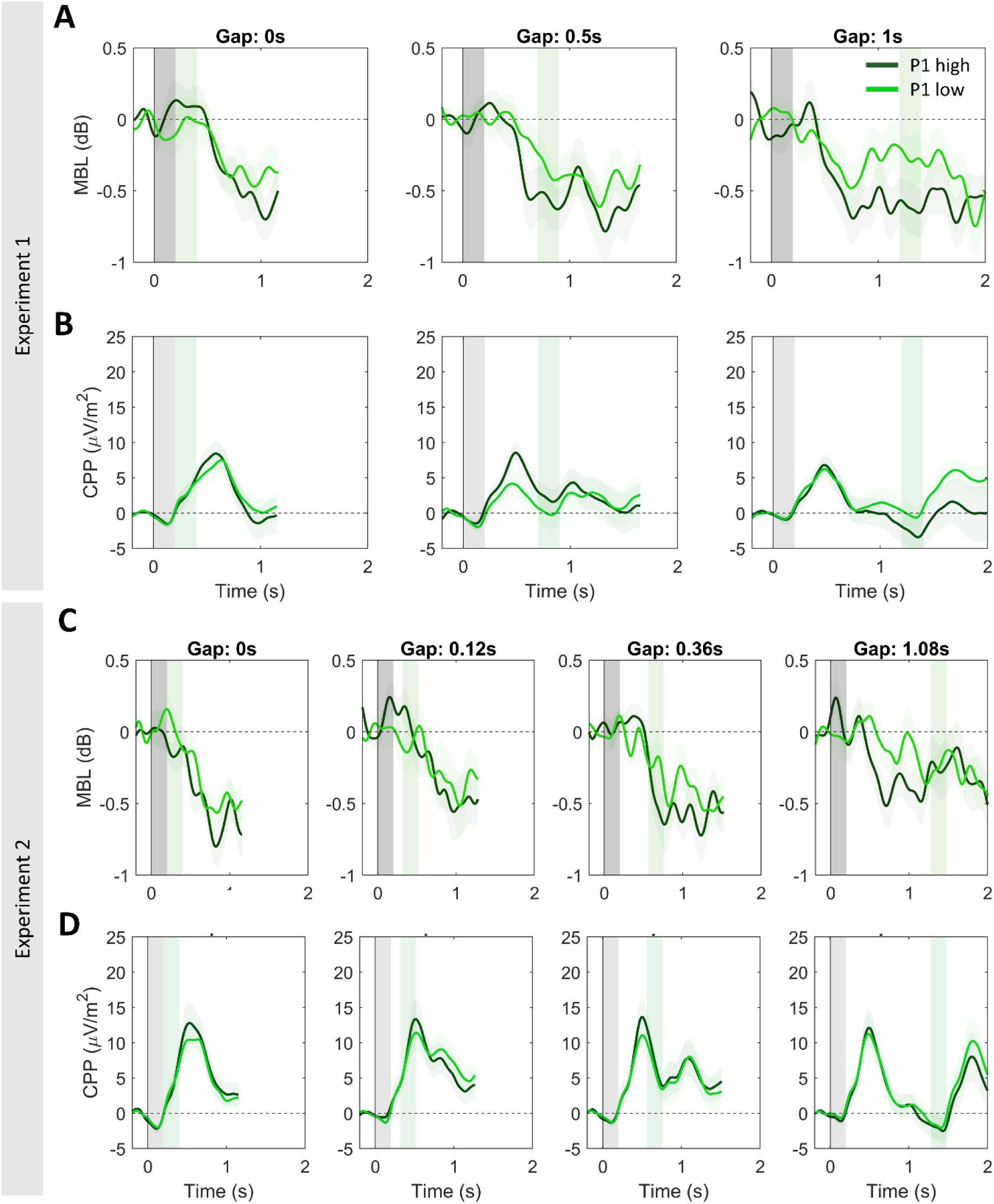
**A,C.** Motor beta lateralisation (MBL) for each gap condition, sorted by P1 coherence in experiments 1 (A) and 2 (C). Lateralisation is computed as a function of actual dot motion [Contra-Ipsilateral hemispheres with respect to correct response hand], sorted by gap duration and P1 coherence pooled across correct & error trials. **B,D.** Centroparietal activity for each gap condition, and sorted P1 coherence in experiments 1 (***B***) and 2 (***D***).

**Figure S4.**
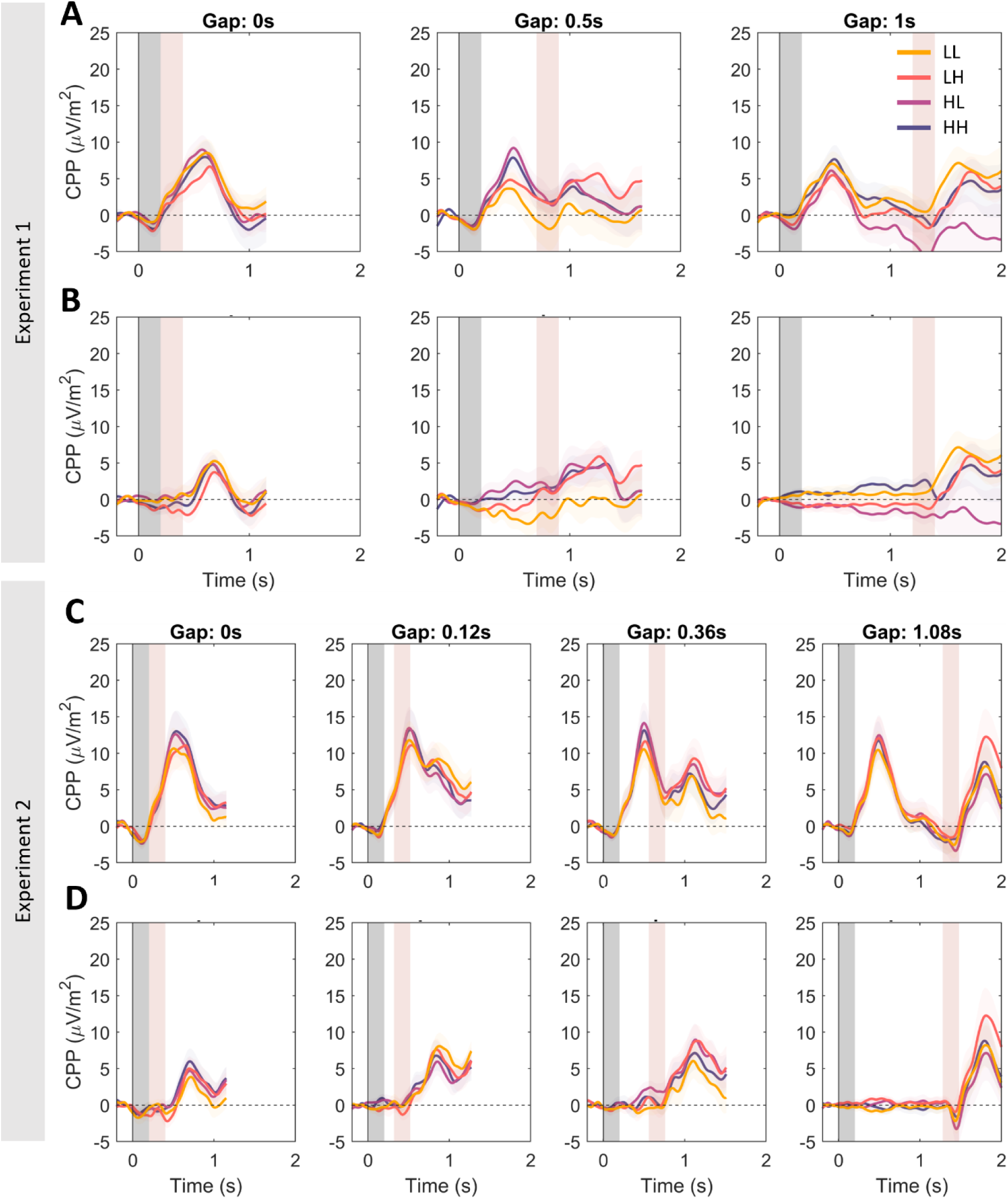
**A,C.** Centroparietal activity sorted by gap, P1 and P2 coherence in Experiments 1 and 2. **B,D.** Centroparietal activity after removing CPP-P1 activity. For each condition, the average of long-gap trials with the same P1 coherence was removed. That is, the average of the long gap HH & HL conditions was subtracted from HH and HL traces. Similarly, the average activity of the long gap LL & LH conditions was subtracted from LL and LH traces. This removes overlapping CPP-P1 potentials, and isolates centroparietal activity related to P2. Subtractions were performed between [-0.2 to 1.4s] from P1 presentation. Thus, CPP-P2 potentials in the long gap condition remain unaltered.

**Figure S5.**
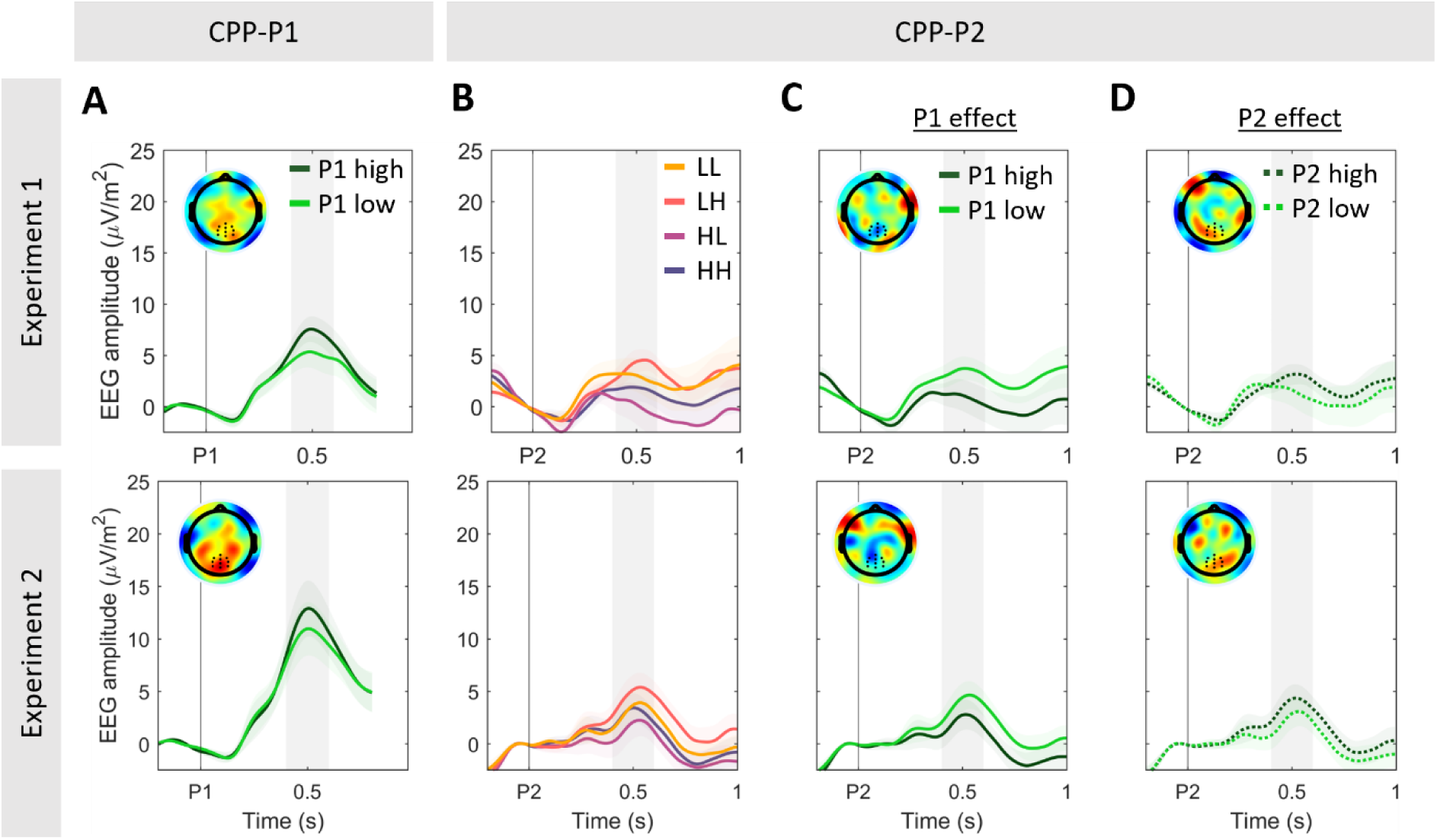
Replication of results in Fig. 3, now performing the analysis on CPP-P2 data (**B-D**) without removing CPP-P1 activity. Note that baseline activity shows a drift consistent with the overlapping potentials from the preceding pulse, which is corrected for when removing CPP-P1 activity (cf. Fig. 3). Note also that the same patterns are qualitatively present in the uncorrected data. Panels C-D illustrate conditions pooled by P1 or P2 coherence, respectively, to highlight the different direction of the effect.

**Figure S6.**
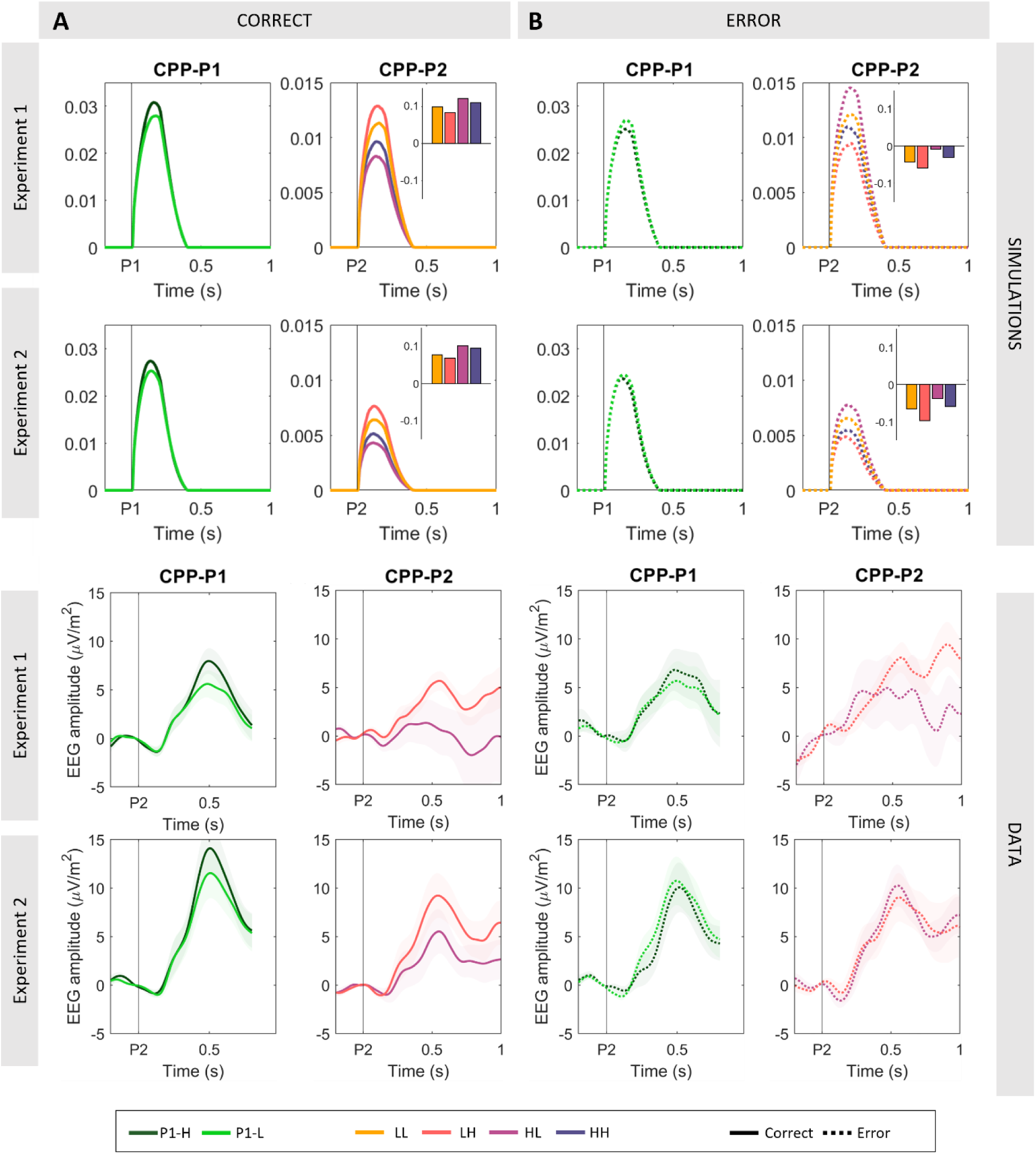
Model-predicted (A-B) and observed (C-D) CPP-P1 and -P2, in correct vs. incorrect responses, for Experiments 1 (top) and 2 (bottom). Bar graph insets in A-B indicate the state of the simulated accumulator at the end of pulse 1 in those trials where the bound was not hit. Positive values indicate that the accumulator was leaning towards the correct answer, whereas negative values indicate that it was leaning towards an error. The model predicts stronger P1 coherence effects in correct compared to error trials (A-B, left), with this effect also being observed in empirical data across both experiments (C-D, left). For the sake of clarity, we do not plot LL and HH conditions in the empirical CPP-P2 data because no clear amplitude differences are predicted by the model in correct vs. error trials, and the trial count in the HH condition was insufficient in most participants. Further, the model predicted a strong interaction in CPP-P2 amplitudes in the HH vs. HL condition. While correct trials (which were the majority of trials in our task and the model) showed higher amplitudes at CPP-P2 for HL compared to the LH condition (A, right), consistent with the effect of P1 on CPP-P1 amplitudes reported in Fig 3, this effect reversed in incorrect trials (B, right). This interaction was also clearly visible in Experiment 2, although slightly less prominent in Experiment 1. Note however that Experiment 1 had fewer overall trials, and power to detect differences is likely to be insufficient after splitting conditions by accuracy.

**Figure S7.**
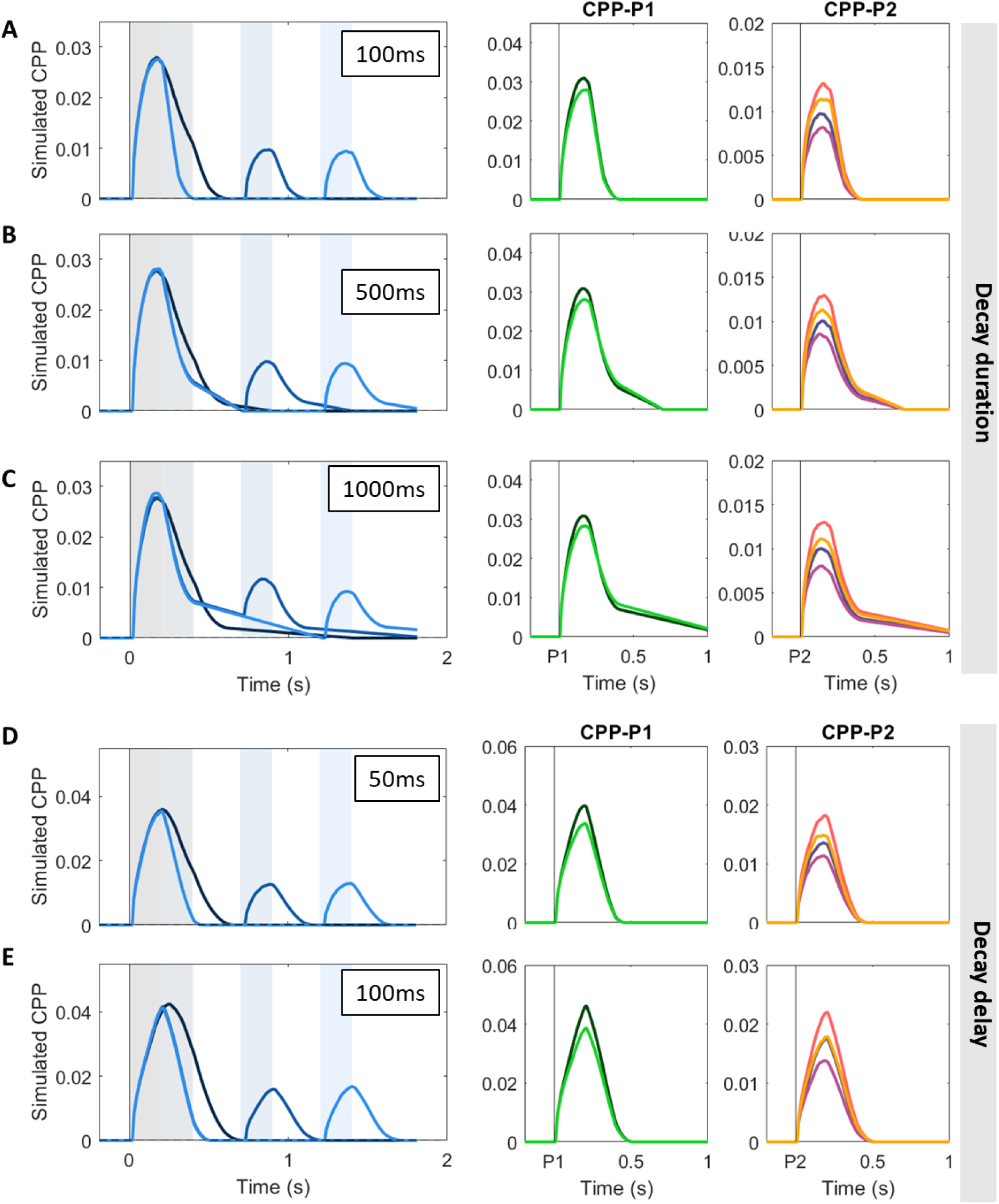
Simulations (N=1000) illustrating the effect of various signal decay time (A-C) and decay delay (D-E) assumptions on the predicted CPP activity. Simulations are based on the fitted model in the main manuscript and for Exp. 1 only for illustration purposes. Given that it’s unclear whether the CPP fall-down to zero when evidence is paused follows the same dynamics as its fall-down when it hits a bound (where studies with immediate behavioural responses suggest that it reaches zero within approximately 200ms; Steinemann et al 2018), we simulated its CPP activity under three different assumptions about the duration of its decay upon evidence interruption (i.e. when blue dots appear onscreen after a pulse). We simulated the CPP assuming the signal takes 100 ms (**A**), 500 ms (**B)** or 1000 ms (**C**) to go back to baseline at pulse offset when a bound had not been reached, retaining a 200 ms fall-down upon bound-crossing. Longer signal decay times for trials where no bound was hit result in a protracted ramp down to baseline. In turn, it is unclear whether any post-decisional evidence accumulation is to be allowed to simulate the CPP in this context, given that previous studies have observed varying delays in signal decay after response (Steinemann et al. 2018; Afacan-Seref et al., 2011). In the manuscript, we assumed no delay between bound crossing and signal decay. Here, we simulated the CPP allowing it to continue building up 50ms (D) or 100ms (E) after bound crossing, while setting it to immediately decay upon blue dot offset. Allowing some post-decisional build-up resulted in higher average CPP amplitudes, and it also better recapitulated some features of the data, including a protracted build-up in the zero-gap condition compared to all others.

**Figure S8.**
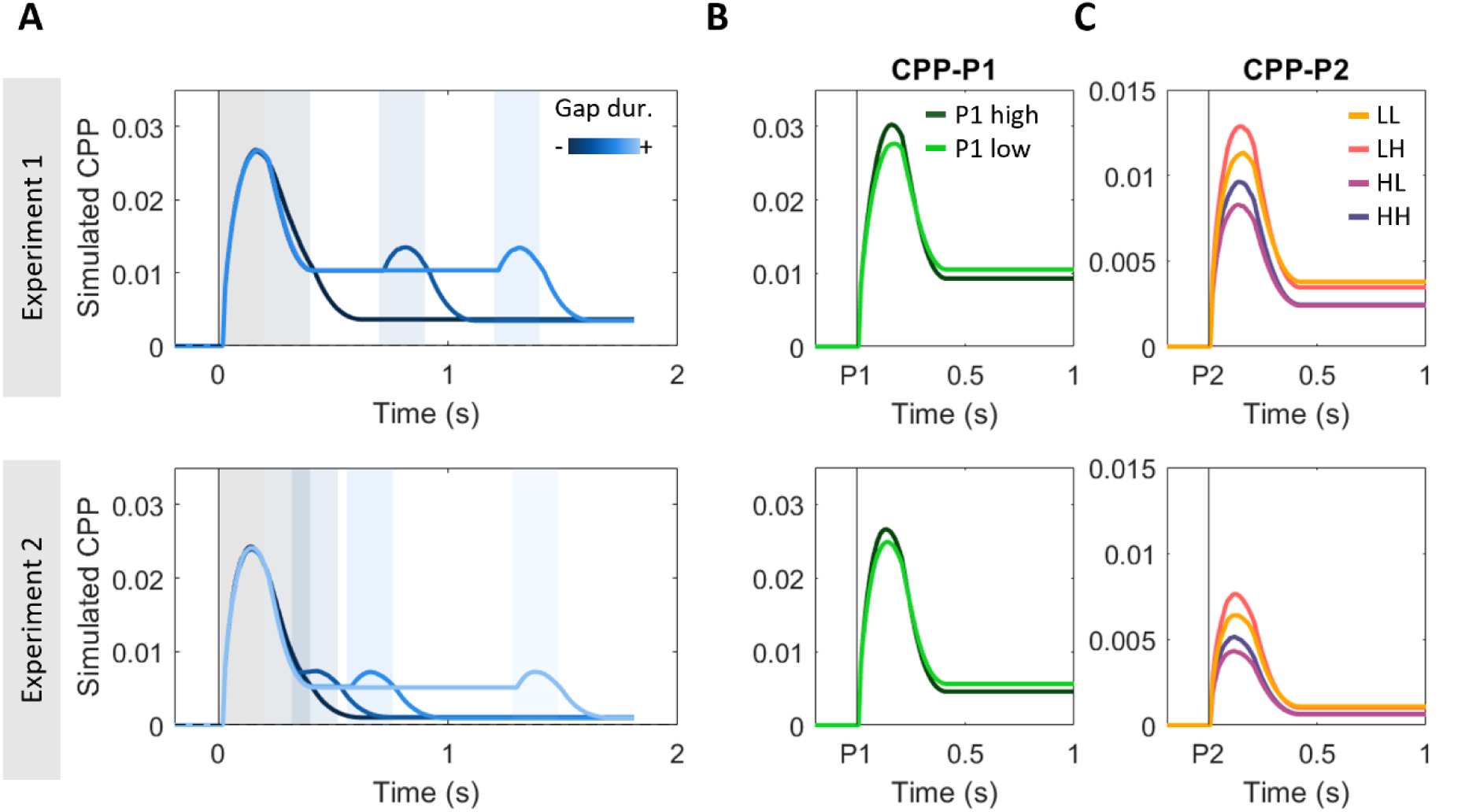
Simulated CPPs using the same computational model as in Fig. 5, but under the assumption that the signal was only reset to baseline if a bound was hit. If the bound was not hit, the accumulator, like the downstream DV at the motor level, was set to stay stable at the final value reached by the end of pulse 1 with no decay and no added noise for the whole duration of the gap. Given that we do not model reaction times, we assumed no transmission delay between events on the screen and their impact on the accumulator. In this model, the partial fall down of the signal during the gap was driven by those trials that hit a bound following the first pulse. Conversely, the fact that it did not fall all the way to 0 on average during the gap is driven by those trials where the DV did not hit a bound during processing of pulse 1 and thus sustained through the gap. Accumulation resumed from that sustained level upon presentation of a second pulse, and the fact that on average CPP-P2 traces reach lower amplitudes than CPP-P1 evoked activities is explained by the fact that trials that were terminated during the first pulse did not engage in accumulation again, and therefore took value 0 in our model. Note that this model provides a qualitatively worse fit to the empirical CPP traces during the gap and CPP-P2 trials. In particular, the magnitude of the CPP-P2 potentials was predicted to be significantly smaller (Exp. 1: CPP-P2 = 19% of CPP-P1 peak amplitude; Exp. 2: CPP-P2 = 13% of CPP-P1 peak amplitude) than observed on the grand-averaged data (CPP-P2 = approximately 50% of CPP-P1 in both experiments; see Fig. 2).

**Figure S9.**
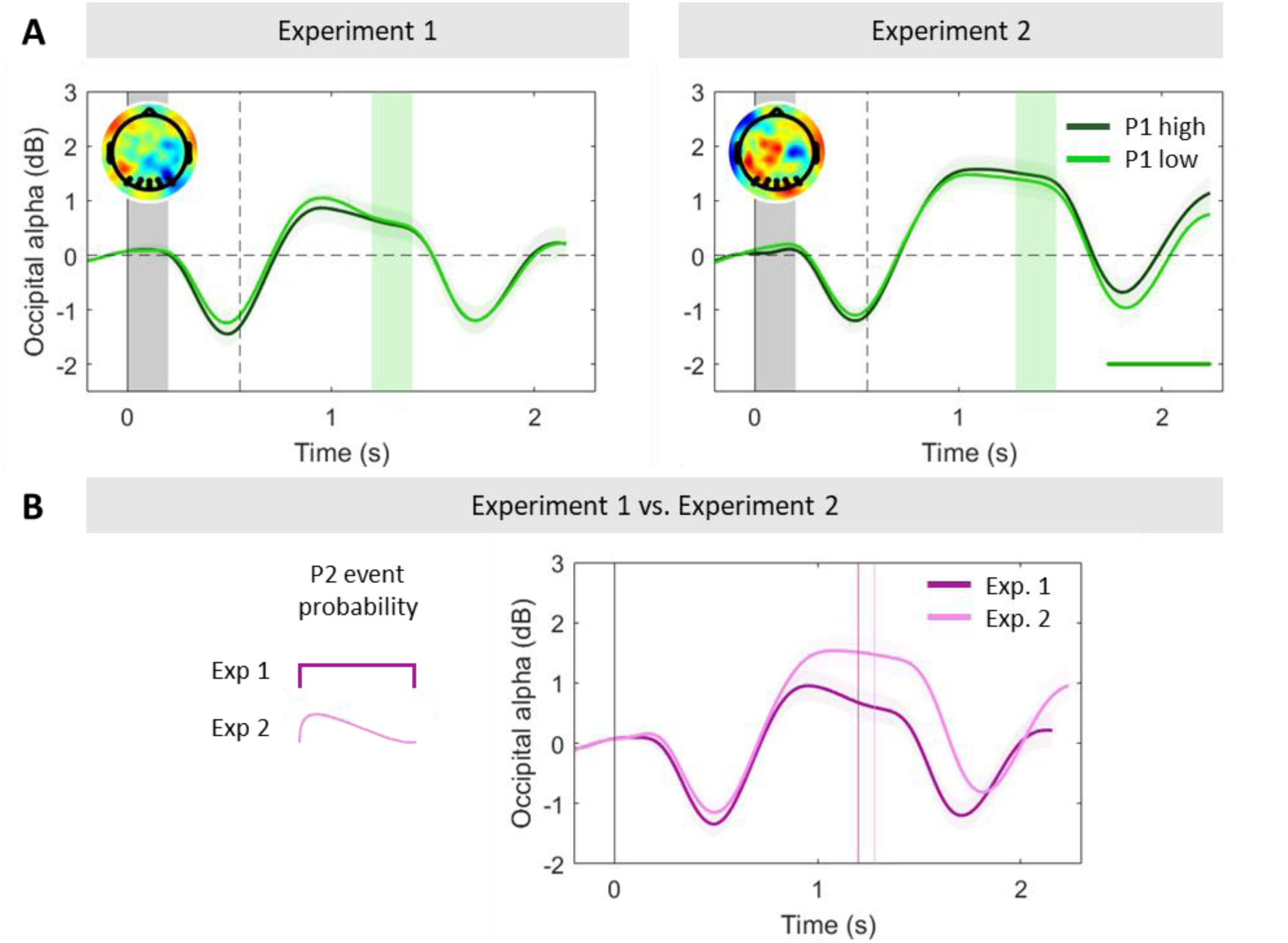
Occipital alpha power tracks dynamics of attentional engagement and varies with temporal expectations. **A.** Grand-averaged (±SEM) occipital alpha power in the long gap conditions sorted by P1 coherence, again baseline-corrected before the onset of the first pulse. At evidence offset, alpha steeply increased, and, interestingly, attained and maintained a higher level during the longer gaps than the baseline before the first pulse (two-tailed cluster permutation test, p < 0.05 in both experiments; see Fig. 2A). One possible interpretation is that this reflects attentional disengagement from the decision occurring after early decision termination. However, if this was the case, we would expect this effect to be dependent on pulse 1 coherence, as it is more likely for a DV to have reached a bound after a high-coherence pulse (Fig. 4B), and we observed no such coherence-dependence in alpha power during the gap. Topographies illustrate the difference in alpha power between P1-high and P1-low coherence trials [-0.2 to 0s] before P2 onset. **B.** Occipital alpha power in the longest gap condition in the two experiments. In Experiment 2, where the event distribution was skewed towards short gap durations between the first and second pulses, participants’ occipital alpha was significantly higher (indicating lower attention) than in Experiment 1 (* p<0.05; two-sample t-test for the difference in alpha power between occipital activity [-0.1 0s] before P2 and the baseline [-0.2 0s] before P1. Alpha power has been linked to temporal expectation encoding (Rohenkohl & Nobre, 2011), as well as being a well-known index of attentional engagement (Thut et al., 2006). Thus, this difference is interesting in light of the difference in gap durations in our two experiments, and hence temporal expectations. In particular, in those trials where no second pulse had been presented after a 0.5s gap, the probability of the second pulse yet appearing was 0.67 in Experiment 1, compared to 0.6 in Experiment 2. Although another difference between experiments was that early terminations within pulse 1 occurred more often in Experiment 2, as predicted by our model (see Fig. 4B), the probability of early terminations is unlikely to be the key influence given the above finding that alpha power during the gap was not dependent on P1 coherence (**A**). Rather than reflecting switches between engagement and disengagement at the decision level, then, it may reflect a gating function at the sensory processing level that operates independent of whether early bound-crossings have happened, yet is modulated between contexts depending on the likelihood of a second pulse still to come. The dotted box indicates the test period. In all panels, shaded areas indicate the timing of evidence pulses for the relevant gap durations. *Markers along the bottom of panel A indicate significant clusters where alpha power differed from zero (two-tailed cluster-based permutation test, p<0.05). In panel B, they indicate significant clusters where alpha power differed in P1-high vs. P1-low coherence trials.

**Figure S10.**
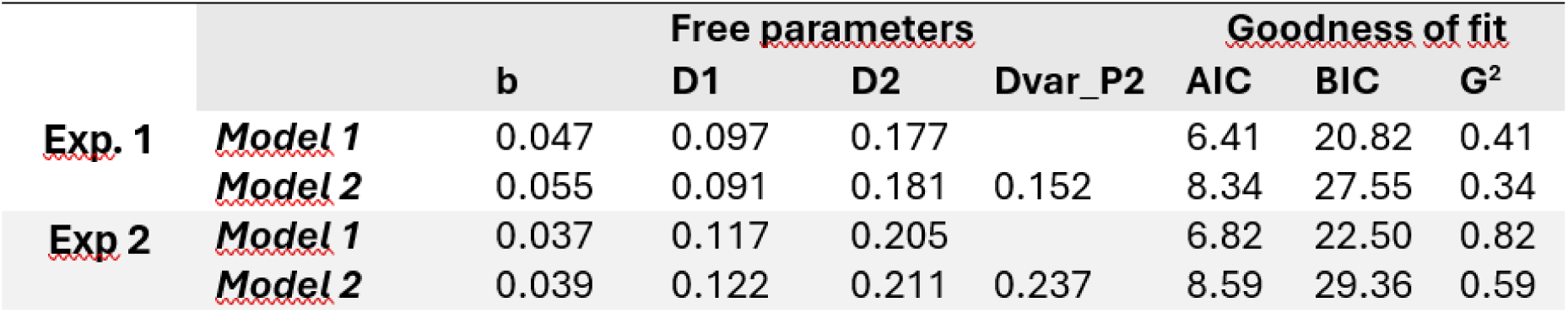
Model comparison investigating the effect of drift rate variability at the second pulse. In our main analysis, we initially fit a model with two different drift rates for low (d1) and high (d2) coherence pulses: **Model 1 (k = 3, b, d1, d2)**. Here, we explored whether increased temporal uncertainty about whether and when a second pulse might appear could have led to more variable attentional states before its appearance, and as a consequence more variable drift rates. To test this hypothesis, we extended the model with one additional free parameter introducing trial-by-trial variability to the drift rates applied to pulse 2: **Model 2 (k = 4: b, d_low, d_high, dvar_p2).** This model resulted in the lowest overall error as measured by G^2^, but the additional complexity was not warranted according to AIC/BIC metrics.

**Figure S11.**
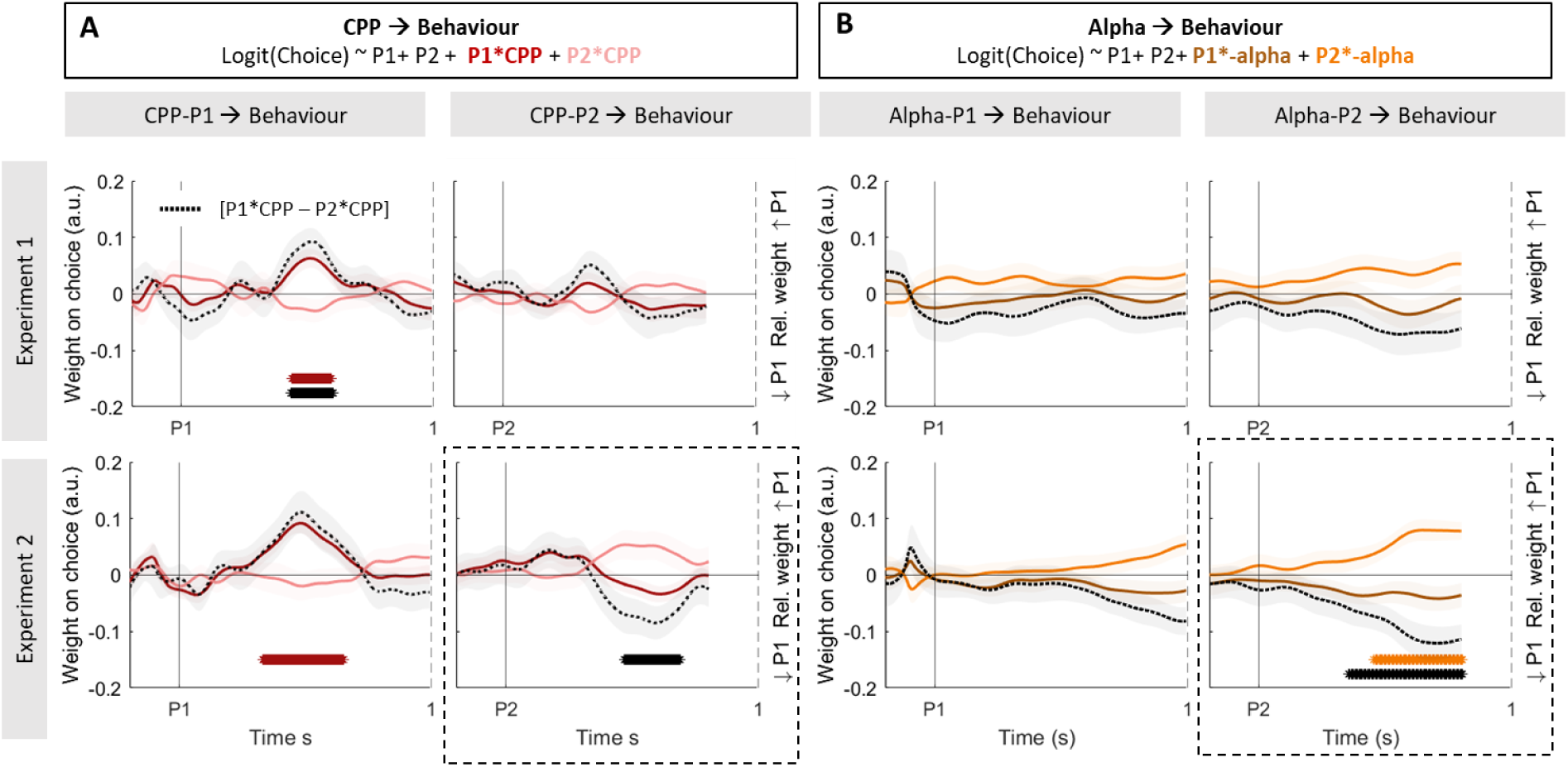
Weight on choice modulations by single-trial fluctuations in neural evidence accumulation and attention markers. In this analysis, we sought to investigate whether single-trial fluctuations in the neural correlates of evidence accumulation and attention were linked to variability in choice behaviour. To do so, we exploited the trial-by-trial variability in the neural signals. **A.** Standardized regression coefficients (±SEM) resulting from Eq. 9, illustrating the effect of CPP-P1 (*left*) and CPP-P2 (*right*) variability on choice, averaged across all gap conditions. The magnitude of CPP-P1 modulated the impact of the first pulse on choices in both experiments while the magnitude of CPP-P2 only modulated the relative weight of P2 in Experiment 2. In our task, variability in the CPP may be related to trial-by-trial variability in motion energy, as well as random noise. Since the weight on choice of P1 and P2 may trade-off with one another due to a bound being set – i.e. on trials where P1 has a strong influence, P2 is likely to have a weaker one - we also computed the difference between the two modulatory terms ([P1coh*CPP - minus P2coh*CPP], for CPP-P1 and CPP-P2 respectively). This difference index effectively quantifies whether centroparietal responses to either pulse modulate the relative weight on choice of each of the pulses. A positive index indicates a relative increase of P1 weight on choice, whereas a negative index indicates a relative increase of P2 weight on choice. Black dashed lines correspond to the difference in regression coefficients for the [P1coh*CPP – P2coh*CPP modulations]. **B.** Standardized regression coefficients (±SEM) resulting from Eq. 10 for alpha aligned to pulse 1 (*left*) or pulse 2 (*right*), averaged across all gap conditions. The magnitude of occipital phasic alpha responses (baselined before P1) did not modulate the weight on choice of P1, and after P2 did modulate its weight on choice in Experiment 2 only. Black dashed lines correspond to the difference in regression coefficients for the P1*alpha – P2*alpha modulations, which effectively indicate a change in the relative weight for P1 and P2. *Asterisks along the bottom of the panels indicate significant (p<0.05) effects in the test period [0.2-0.8s post event].

**Figure S12.**
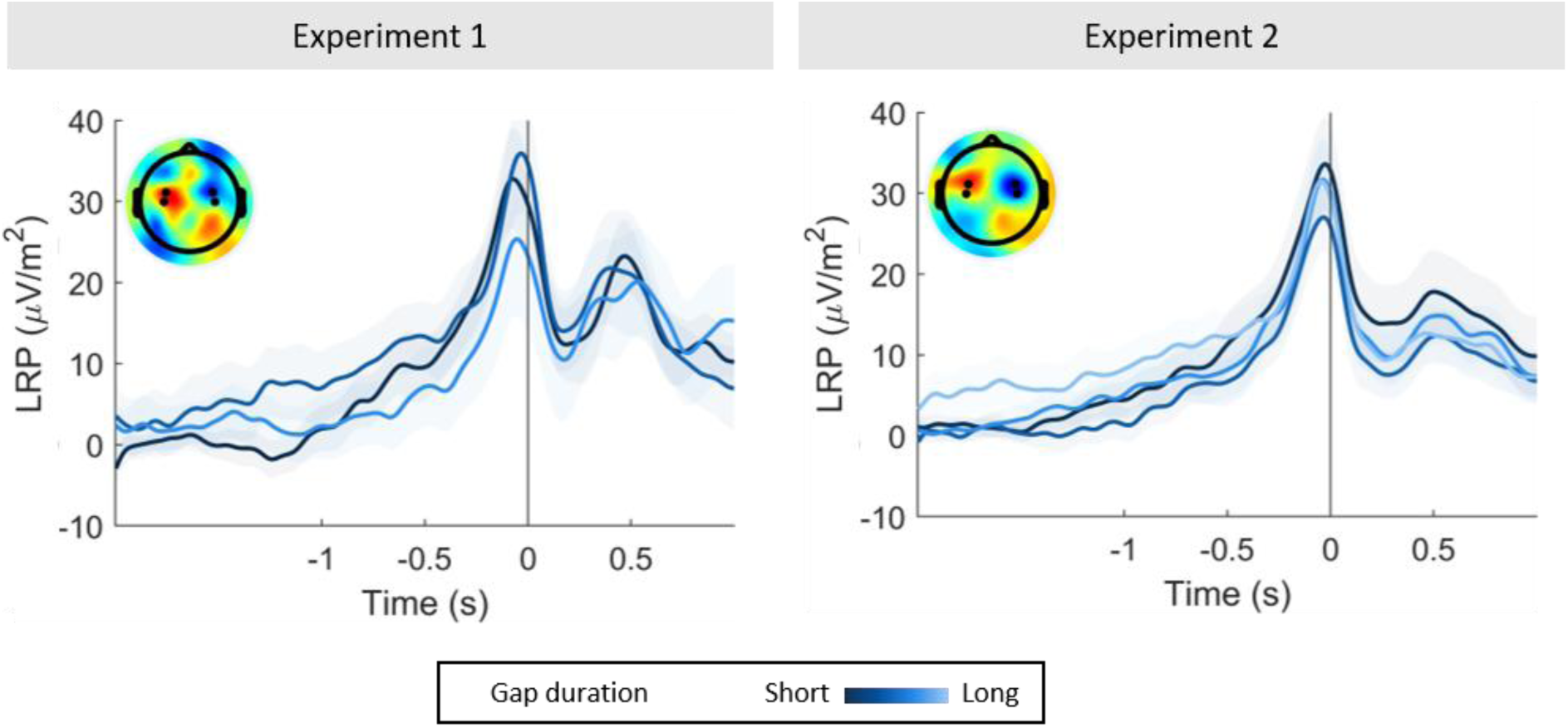
Grand-averaged (±SEM) lateralised readiness potential (LRP), computed as the difference in voltage between [Ipsi - Contralateral] hemispheres with respct to the correct response hand, and sorted by gap duration, in correct trials only. Higher LRP values indicate stronger negativities over the contralateral hemisphere. Topography insets indicate the difference in voltage across the scalp for [Left - Right]-hand responses [-0.1s to 0s] before action.

**Figure S13.**
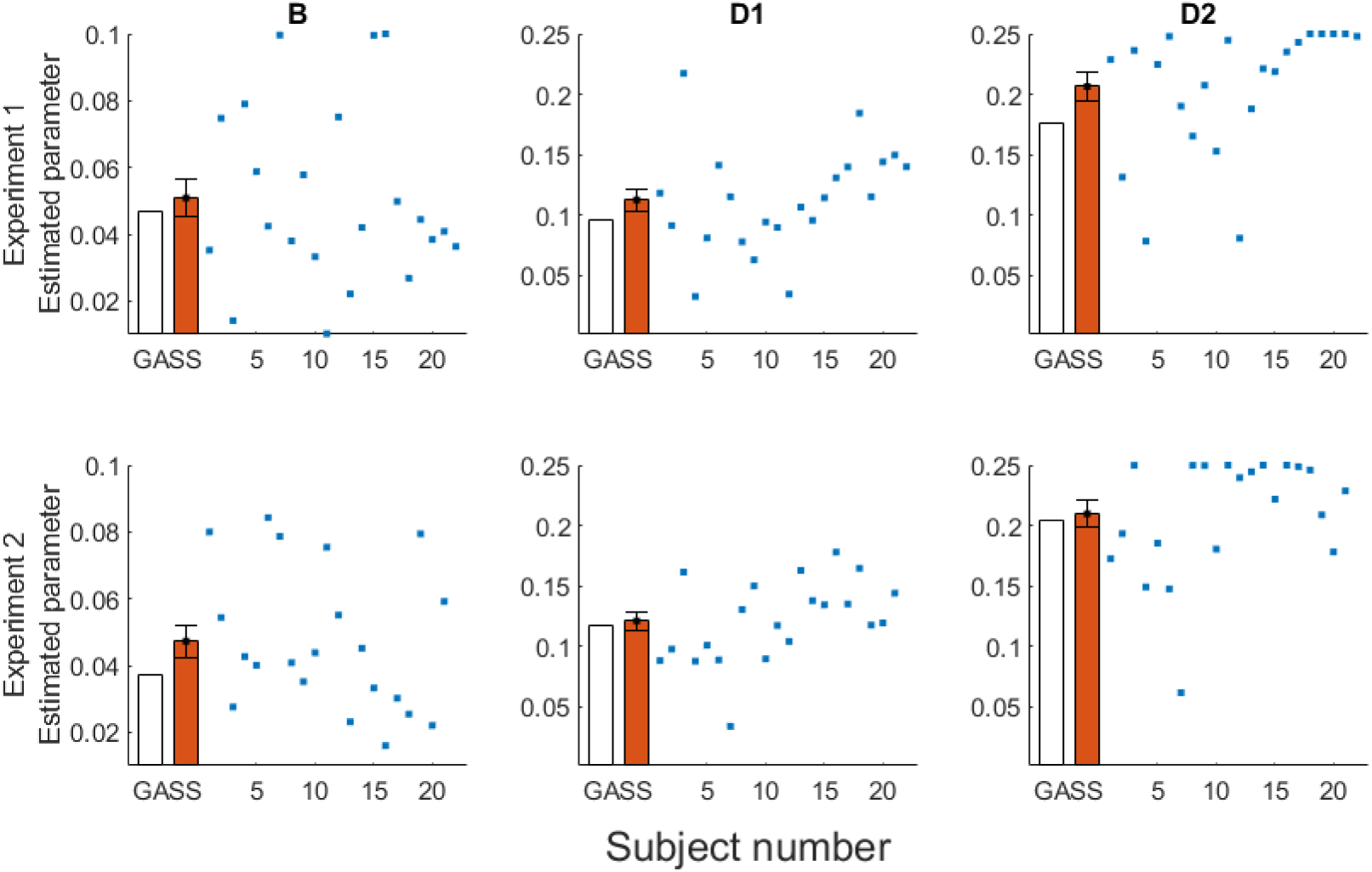
Model parameters (bound (B), low drift rate (D1), high drift rate (D2)) obtained using grand-averaged data (white bars) in each experiment, used for the analyses and simulations in the main manuscript. To assess the robustness and variability of these results across participants, we additionally fit the same model to each individual participant (N = 5000, blue dots). Orange bars indicate the mean (± SEM) of these individually-estimated parameters. The averaged parameters qualitatively recapitulate the results from the model based on grand-averaged data.

**Figure S14.**
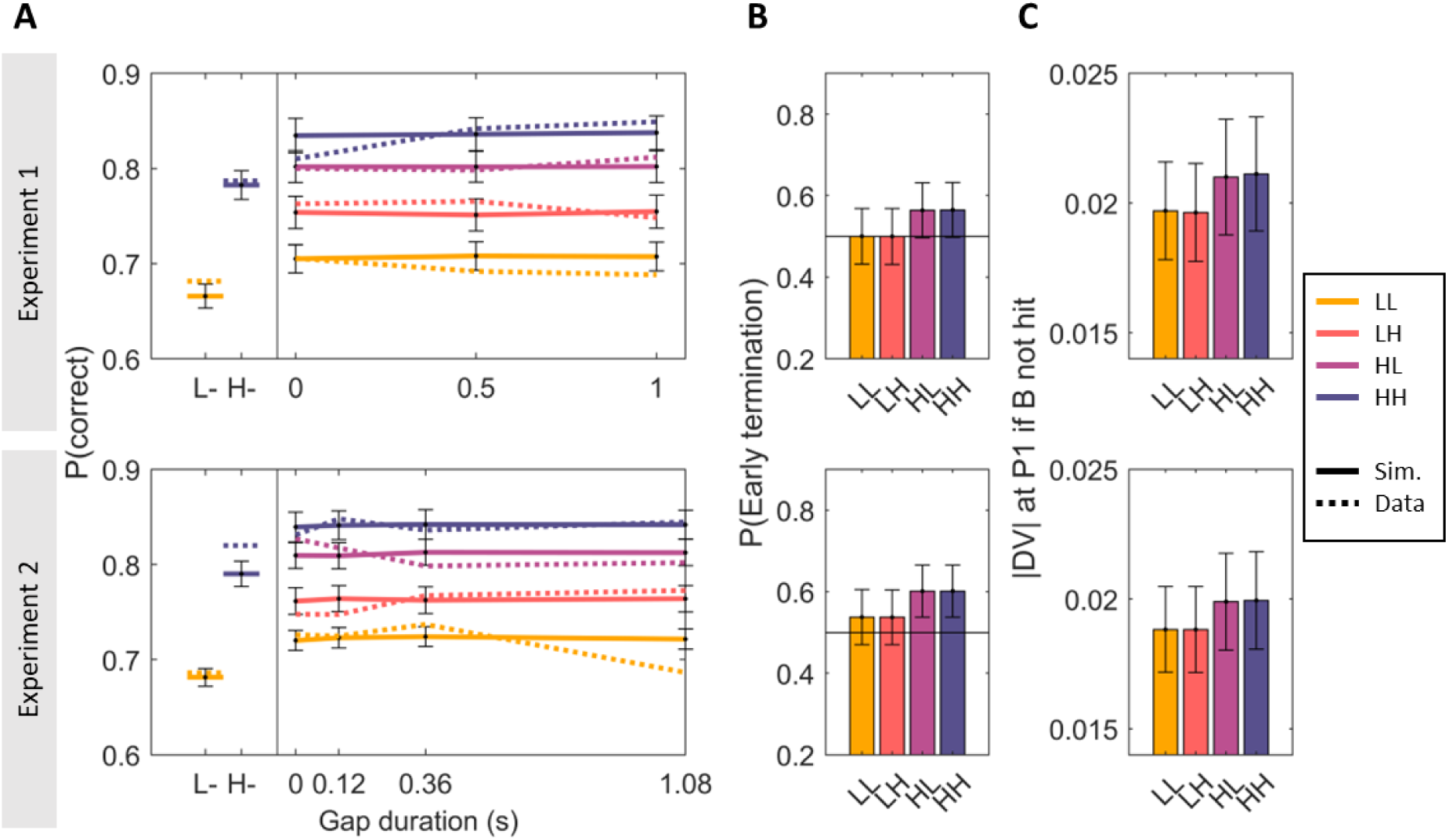
Replication of results in Figure 4, using models fit to individual participants rather than to the grand-averaged data. We used the individually-fit parameters (B (bound), d1 (low drift rate), d2 (high drift rate)), to simulate N = 10000 for each participant. Data shown here represent the average of all simulations across participants, and error bars indicate the Standard Error of the Mean (SEM). These simulations of individual data fits closely match the results obtained using the model fit to the grand-averaged data. **A.** Grand-averaged simulated (*solid lines*) accuracy in single-(*left*) and two-pulse (*right*) trials, sorted by pulse coherence and gap duration in experiments 1 (*top*) and 2 (*bottom*). Observed data are overlaid (*dashed*) for comparison. **B.** Proportion (mean ± SEM) of simulated two-pulse trials that hit a bound during the first pulse, and therefore did not use the second pulse evidence to make a choice. **C.** Absolute value (mean ± SEM) of the decision variable (DV) at the end of pulse 1 in the subset of trials where the bound was not hit during that pulse, sorted by condition.

## Supplementary Note 1

We explored alternative models that could explain both the key behavioural and neural patterns of results in our dataset, and compared both the fit quality to accuracy data and the predicted EEG results to our original implementation. All models were fit to the grand-averaged accuracy data for all conditions, using 10K simulations, and for both experiments separately (as was done for the bounded model presented in the manuscript). For EEG simulations in unbounded models, we assumed that the CPP signal fell back down to zero upon the dots turning blue, as for the simulations in the main manuscript. **Fig. S15** below illustrates the fit quality as measured by means of Bayesian Information Criterion (BIC).

**Fig. S15.**
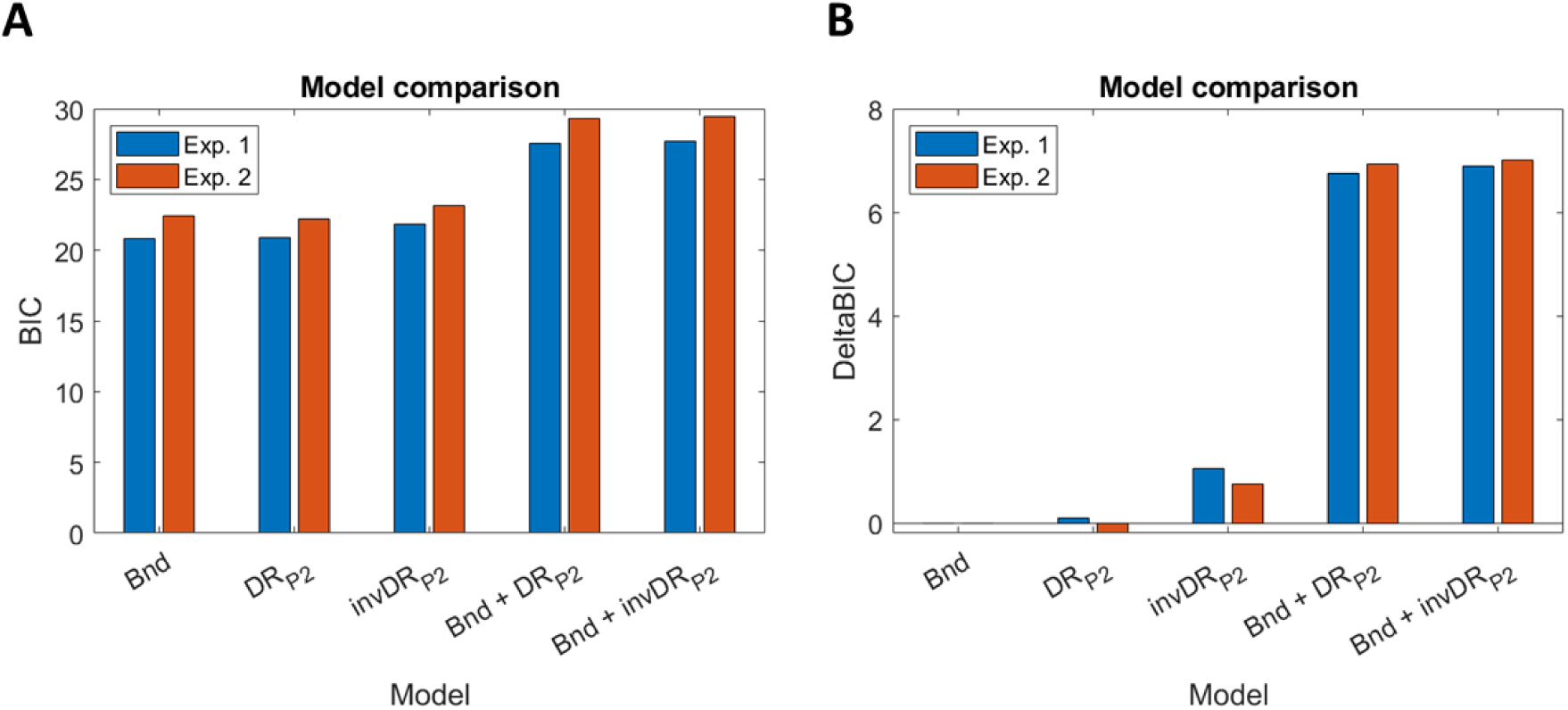
Model comparison. **A-B.** Absolute (A) Bayesian Information Criterion (BIC) for each model, or relative (B) to the bound-only model presented in the original paper. The compared models include the original bounded model implementation in the manuscript (Bnd); an equally complex model with no bound and a P2 drift rate scaling parameter (DR_P2_); a model with no bound, but a parameter that scaled P2 drift rates inversely to P1 coherence (invDR_P2_); an extension of the original model containing both a bound and a P2 drift rate scaling parameter that systematically downweighted P2 evidence (B + DR_P2_) or downweighted P2 evidence in a manner inversely proportional to P1 coherence (B + invDR_P2_).

### 1. Unbounded model + P1-independent P2 drift rate reweighting (*DR_P2_*)

First, we fit a model with no bound in which the drift rate for P2 was estimated by reweighting (positive or negative) the P1 drift rate via an additional free scaling parameter *w*. This model thus had the same complexity as our original one (k = 3 free parameters): two drift rate parameters for high and low coherence of P1 (*d_high_*, *d_low_),* plus a scaling parameter, *w*, dictating the strength of P2 relative to same-coherence P1. The drift rate for P2 (d_P2_) was computed as the d*w, where d equals d_high_ or d_low_ depending on P2 coherence, multiplied by the scaling parameter *w,* so that if w < 1, P2 drift rates would decrease compared to the same coherence in P1.

This first model (DR_P2_) yielded a similar fit quality to the data compared to our original bounded model (Bnd), as measured by BIC (**Fig. S15**). The new model broadly recapitulated the key behavioural results, including order effects, (**Fig. S16,A**) and could also recapitulate the generally lower CPP-P2 amplitudes (**Fig. S16,B**). However, this model failed to capture the key coherence-based pattern observed in the CPP-P2 data. Namely, while our EEG results showed that CPP-P2 in trials following P1-low coherence pulses reached overall higher amplitudes than that in trials following P1-high coherence pulses (see Fig. 3), this model’s simulations predicted that CPP-P2 should scale only with P2 coherence, showing higher amplitudes and steeper build—up rates for P2-high trials, regardless of P1 coherence (**Fig. S16, C**). This is at odds with our empirical results.

**Fig. S16.**
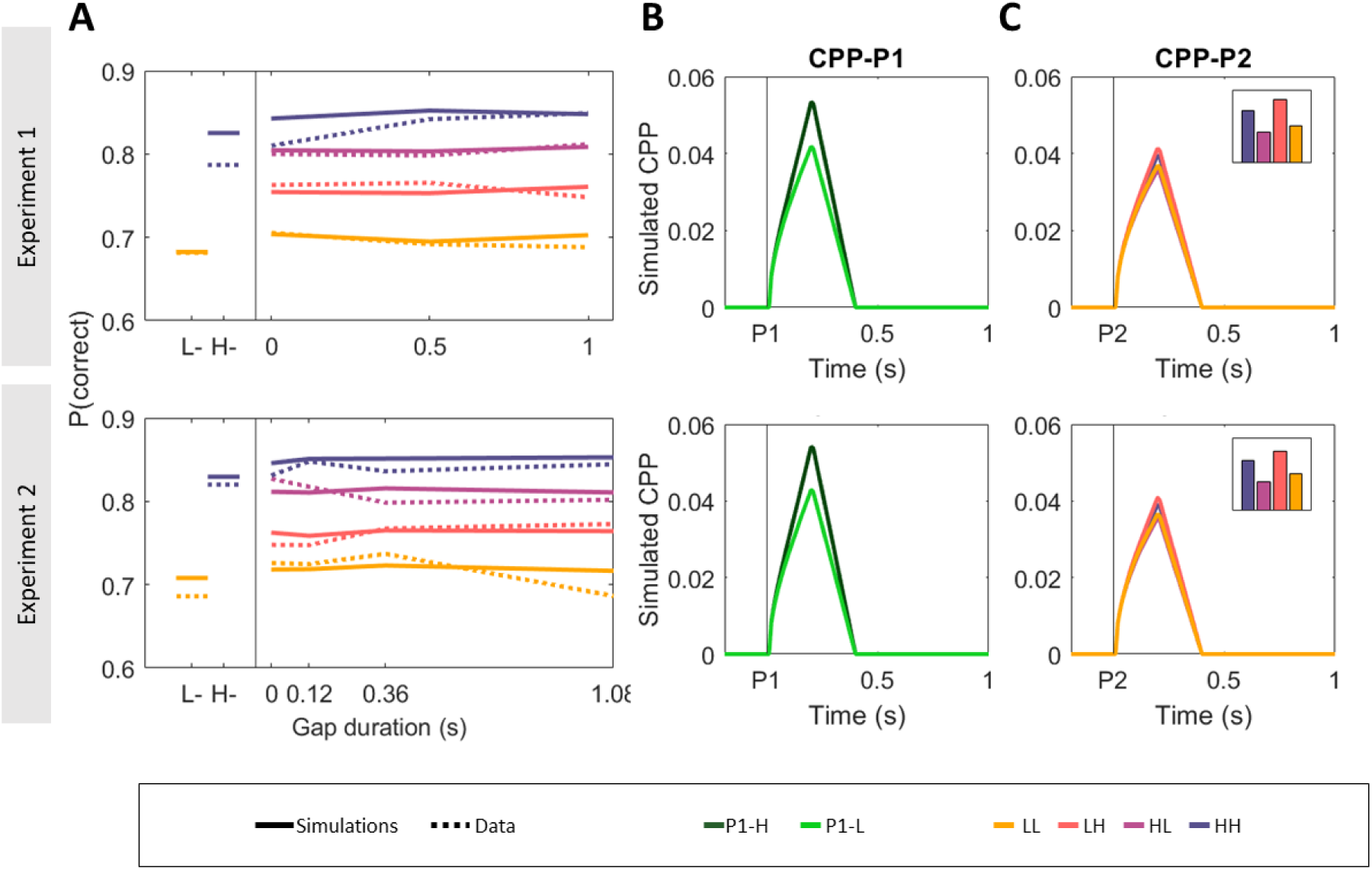
Simulation results (N = 10K) of a model with no bound, and a free parameter that systematically reweighted P2 drift rates relative to P1. **A.** Observed (*dashed*) and simulated (*solid*) proportion of correct responses. **B.** Simulated CPP-P1 traces, sorted by P1 coherence. **C.** Simulated CPP-P2 coherences, sorted by P1 and P2 coherence. Bar graph insets indicate average CPP-P2 amplitude [0-0.2s after P2] for visualisation purposes. High-coherence P2 trials are predicted to reach higher amplitudes than low-coherence P2 ones, irrespective of P1 coherence. Bar graph insets indicate average CPP-P2 amplitude [0-0.2s after P2]. This is at odds with our empirical findings (c.f. Fig. 3 in manuscript).

### 2. Unbounded model + P1-dependent P2 Drift rate reweighting (invDRP2)

Next, we tested a model in which P2 downweighting could depend on P1 strength. That is, we made the P2 drift rate scaling parameter *w* inversely proportional to P1 coherence so that P2 drift rate d_P2_ = d*w/d_P1_, where d equals d_high_ or d_low_ depending on P2 coherence, and d_P1_ indicates the preceding P1 coherence. This implements a kind of certainty weighting, whereby evidence following a strong P1 is more strongly dampened than evidence following a weak P1. This model could recapitulate the key behavioural findings (**Fig. S17,A**), and also qualitatively captured the CPP-P2 effects (i.e. P1-low trials reaching overall higher amplitudes than P1-high trials, **Fig. S17,C**), although the magnitude of this effect was substantially smaller than predicted by the original simple bounded model with no drift rate modulations. However, BICs indicated that this model provided an overall worse fit to the accuracy data compared to the original bounded model (**Fig. S15**).

**Fig. S17.**
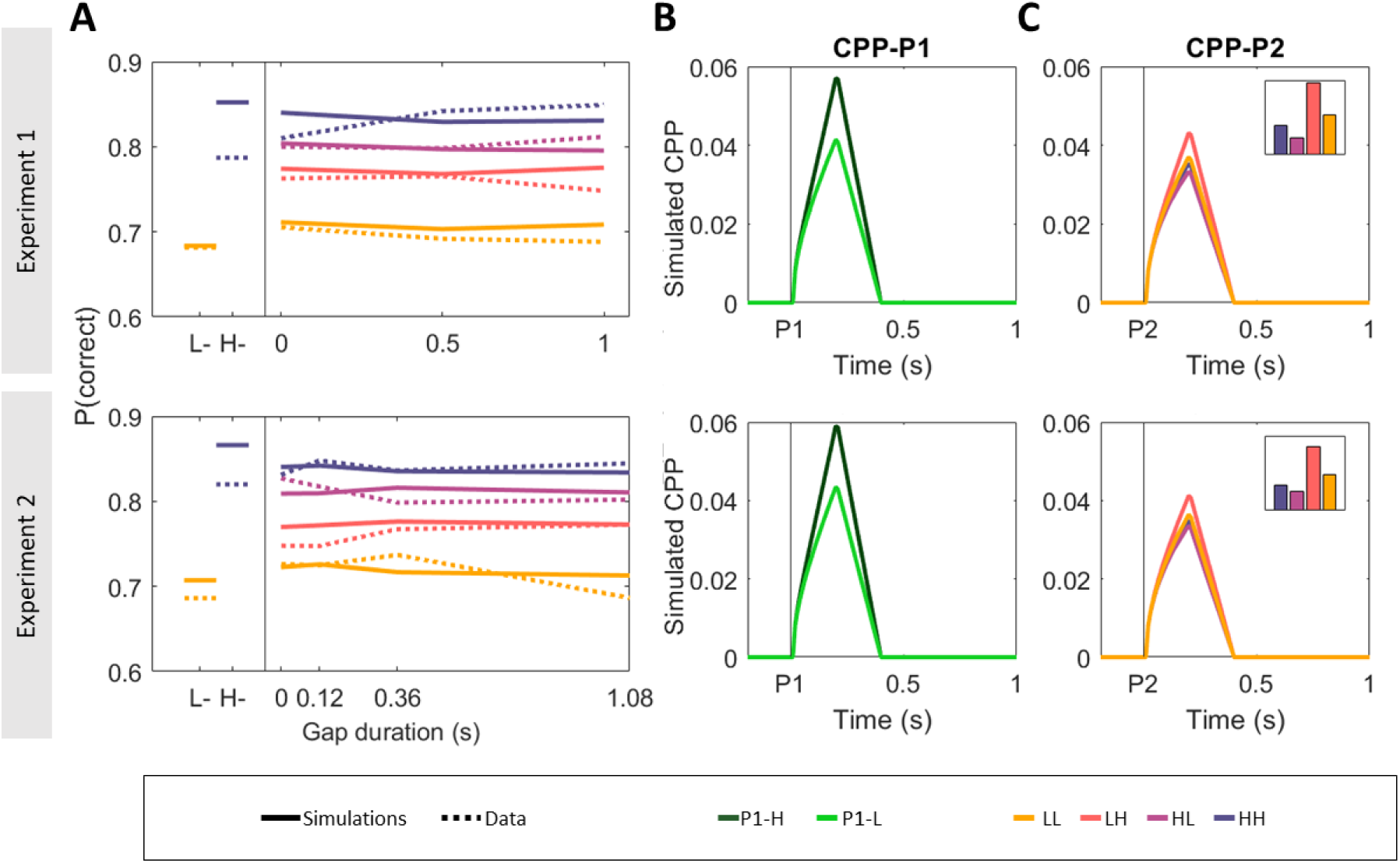
Simulation results (N = 10K) of a model with no bound, and a free parameter that scaled P2 drift rates in a manner inversely proportional to P1 coherence. **A.** Observed (*dashed*) and simulated (*solid*) proportion of correct responses. **B.** Simulated CPP-P1 traces, sorted by P1 coherence. **C.** Simulated CPP-P2 traces, sorted by P1 and P2 coherence. Bar graph insets indicate average CPP-P2 amplitude [0-0.2s after P2] for visualisation purposes. CPP-P2 amplitudes scaled with both P1 and P2 coherences, in line with our empirical findings, though the fit to behaviour was compromised (see BICs in Fig S15).

### 3. Bounded models + P2 drift rate reweighting (*Bnd + DR_P2,_*, *Bnd+ invDR_P2_*)

Finally, we investigated how well models with both a bound and either of the two P2 drift rate scaling methods we investigated above (uniform downweighting, P1-dependent downweighting) could capture the data, thus effectively testing two extensions of our original implementation that allowed for flexible reweighting of P2.

The uniform downweighting model with a bound (Bnd + DR_P2_) could recapitulate all key behavioural and EEG findings (**Fig. S18**, but the additional complexity of the model was not supported by BIC (**Fig. S15**). Crucially, the reason that this model could recapitulate the CPP-P2 results was still the presence of a bound, although we note that the proportion of trials that were predicted to terminate early was reduced in this model compared to the original bounded model presented in the manuscript (c.f. Fig. 4). Yet, this relatively small fraction of trials where accumulation ended early meant that 1) in some trials no accumulation was allowed to occur at all during P2, and 2) where it occurred, the DV was closer to bound following P1-high coherence trials, thus needing to accumulate less further evidence before reaching a bound and yielding smaller CPP-P2 amplitudes overall in those trials. The inclusion of a bound was thus key to allow a model with uniform P2 downweighting to account for both behavioural and EEG data.

The model with P1-based scaling of P2 drift rates with a bound could also recapitulate all key behavioural and EEG findings (**Fig. S19**), but again, the additional complexity of the model was not supported by BICs (**Fig. S15**).

**Fig. S18.**
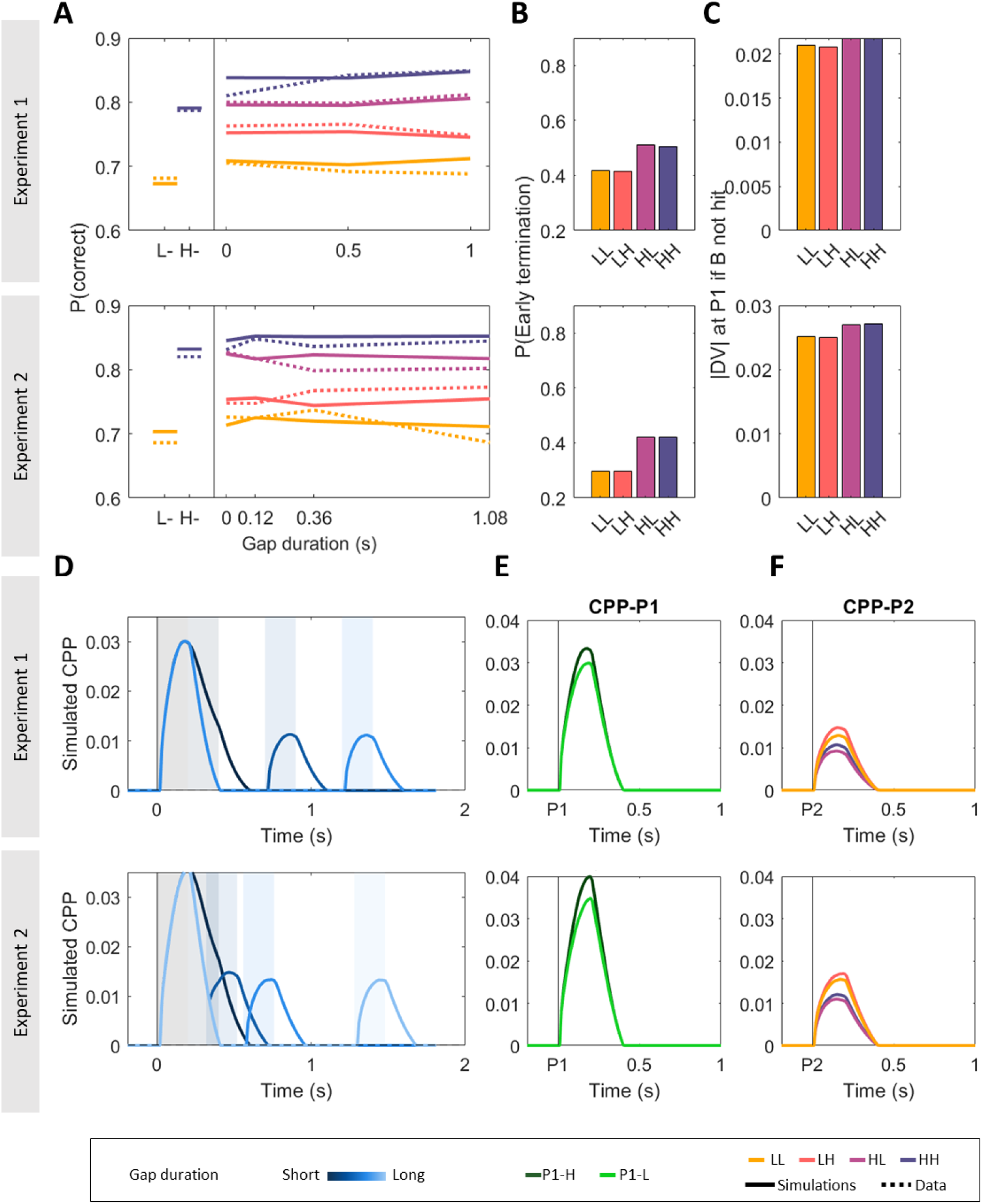
Simulation results (N = 10K) of an extension of our original model with a bound, two drift rates, and an additional free parameter that systematically scales P2 drift rates (Bnd + DR_P2_), irrespective of P1 coherence. This model could recapitulate all key behavioural and EEG findings. **A.** Observed (*dashed*) and simulated (*solid*) proportion of correct responses. **B.** Model-estimated proportion of trials that terminated during pulse 1. **C.** Estimated absolute value of the decision variable at the end of P1, if no bound was hit. **D-F.** Model-based CPP simulations sorted by gap duration (**D**), P1 coherence and aligned to P1 (**E**), or P1-P2 coherence and aligned to P2 (**F**).

**Fig. S19.**
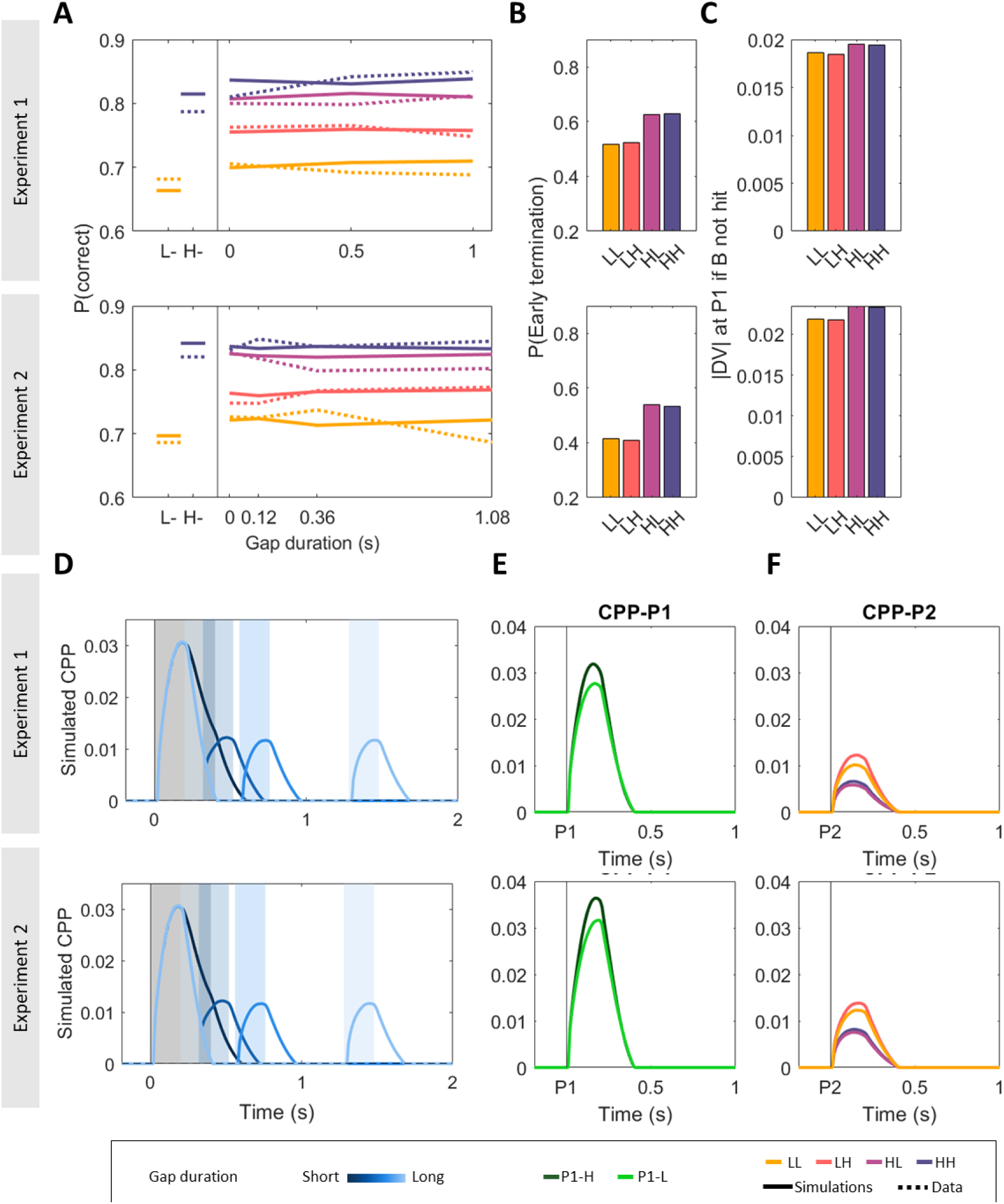
Simulation results (N = 10K) of an extension of our original model with a bound, two drift rates, and an additional free parameter *w* that inversely scales P2 drift rates with P1 coherence (Bnd + invDR_P2_), so that dP2 = d*w/dP1. This model could recapitulate all key behavioural and EEG findings. **A.** Observed and simulated proportion of correct responses. **B.** Model-estimated proportion of trials that terminated during pulse 1. **C.** Estimated absolute value of the decision variable at the end of P1, if no bound was hit. **D-F.** Model-based CPP simulations sorted by gap duration (**D**), p1 coherence and aligned to P1 (**E**), or P1-P2 coherence and aligned to P2 (**F**).

## Conclusion

This extended modelling exercise suggests that 1) a simple bounded model is favoured by model comparison, 2) an alternative model of equal complexity which includes a P1-dependent systematic downweighting of P2 rather than a bound can produce qualitatively similar results, the common feature of both viable models being the push-pull relationship between P1 and P2, and 3) more complex models including both a bound and P2 modulations can also account for both the behavioural and EEG results, but the additional complexity is not supported by the current data. While it is possible that some flexible weight modulations occur, these are not sufficiently influential to justify its inclusion in the model. Future work would nevertheless be warranted to explore this possibility in more detail using tailored task paradigms.

